# Comparative genomics of emerging lineages and mobile resistomes of contemporary broiler strains of *Salmonella* Infantis and *E. coli*

**DOI:** 10.1101/2021.01.22.427796

**Authors:** Ama Szmolka, Haleluya Wami, Ulrich Dobrindt

## Abstract

**Introduction:** Commensal and pathogenic strains of MDR *Escherichia coli* and non-typhoid strains of *Salmonella* represent a growing foodborne threat from foods of poultry origin. MDR strains of *S*. Infantis and *E. coli* are frequently isolated from broiler chicks and the simultaneous presence of these two enteric bacterial species would potentially allow the exchange of mobile resistance determinants.

**Objectives:** In order to understand possible genomic relations and to obtain a first insight into the potential interplay of resistance genes between enteric bacteria, we compared genomic diversity and mobile resistomes of *S*. Infantis and *E. coli* from broiler sources.

**Results:** The cgMLST analysis of 56 *S*. Infantis and 90 *E. coli* contemporary strains revealed a high genomic heterogeneity of broiler *E. coli*. It also allowed the first insight into the genomic diversity of the MDR clone B2 of *S*. Infantis, which is endemic in Hungary. We also identified new MDR lineages for *S*. Infantis (ST7081 and ST7082) and for *E. coli* (ST8702 and ST10088). Comparative analysis of antibiotic resistance genes and plasmid types revealed a relatively narrow interface between the mobile resistomes of *E. coli* and *S*. Infantis. The mobile resistance genes *tet(A), aadA1* and *sul1* were identified at an overall high prevalence in both species. This gene association is characteristic to the plasmid pSI54/04 of the epidemic clone B2 of *S*. Infantis. Simultaneous presence of these genes and of IncI and IncX plasmids in cohabitant caecal strains of *E. coli* and *S*. Infantis suggests an important role of these plasmids in a possible interplay of resistance genes between *S*. Infantis and *E. coli* in broilers.

**Conclusions:** This is the first comparative genomic analysis of contemporary broiler strains of *S*. Infantis and *E. coli*. The diversity of mobile resistomes suggests that commensal *E. coli* could be potential reservoirs of resistance for *S*. Infantis, but so far only a few plasmid types and mobile resistance genes could be considered as potentially exchangeable between these two species. Among these, IncI plasmids could make the greatest contribution to the microevolution and genetic interaction between *E. coli* and *S*. Infantis.

## 1. Introduction

Reoccuring emergence of multidrug-resistant (MDR) strains of *Salmonella enterica* serovar Infantis is a global challenge for public health and food safety. Although it is accepted that the use of antibiotics in animal industry is one of the major driving forces of this process, there are several factors influencing development and emergence of MDR non-typhoidal *Salmonella* in food-producing animals. In most industrialized countries the poultry (primarily layer and broiler as well as fattening turkey) is the major source of foodborne salmonellosis. During the last two decades, MDR clones of *S*. Infantis have emerged and became established in the broiler industries of several countries in Europe (EFSA 2020), in Israel, Turkey, Japan and in the USA (Shahada et al, 2006; Gal-Mor, et al., 2010; Tate et al., 2017; Acar et al, 2019) respectively.

In general, enteric bacteria such as *E. coli* and *Salmonella* are readily responding to the frequent use of antibiotics by developing resistance. Due to their great genomic plasticity, *E. coli* strains can be more adaptive to the unfavourable antibiotic environment (Mathew et al., 2002). Part of this flexibility is the ability to acquire resistance determinants by horizontal gene transfer, and to develop new MDR lineages by vertical gene transfer ensuring survival and persistence. Based on this difference between *Salmonella* and *E. coli*, it is logical to assume that the resistome of *E. coli* in animals may serve as a reservoir for drug resistance determinants for *Salmonella* (Szmolka and Nagy, 2013). This could be especially true for the broiler industry where the high stocking density and the frequently necessitated antimicrobial treatments could fuel the mobilization and uptake of resistance determinants within the intestinal *E. coli* population and between *E. coli* and *Salmonella* (Gyles, 2008). Indeed, some studies revealed *in vivo* or semi *in vivo* AMR gene transfers between *Salmonella* and *E. coli* in chicks (Oladeinde et al., 2019) and in turkey poults (Poppe et al., 2005) or in simulated intestinal environment of pigs (Blake et al., 2003) with or without simultaneous application of antibiotics.

Regarding the growing economic and public health importance of MDR *Salmonella* Infantis and the situation of MDR *E. coli* of broilers described above, it seems to be logical to ask how much the mobile resistomes of these two important MDR enterobacterial species overlap in real life as an indication of possible exchanges of resistance genes. So far, only a few studies compared the resistance phenotype and genotype of contemporary *Salmonella spp*. and *E. coli* isolates from poultry (Varga et al., 2019) and from broilers (Ahmed et al., 2009), but they have been mainly carried out on a phenotypic basis or have only investigated selected resistance genes of a relatively low number of strains.

Therefore, we performed a comparative resistome analysis at the whole-genome level of contemporary of *S*. Infantis isolates from humans and broilers and of commensal and extraintestinal pathogenic *E. coli* (ExPEC) strains isolated from broilers. With these studies, we aimed to gain a first insight into the possible interspecies exchange of mobile resistance determinants between *E. coli* and *S*. Infantis circulating in the broiler industry. Our findings indicated that certain plasmid types and resistance genes or gene cassettes could be considered in a possible genomic interaction between *E. coli* and *S*. Infantis.

## 2. Materials and methods

### 2.1. Sampling of *S*. Infantis and *E. coli* isolates

Whole-genome sequences of 56 *S*. Infantis and 90 *E. coli* isolates were analysed in this study. The collection was established in order to provide a genome-based comparison of antibiotic resistance in *S*. Infantis and *E. coli* from different broiler sources in Hungary.

*S*. Infantis strains were mostly isolated from the faeces of broiler chickens (n=31) and from human samples (n=25) representing sporadic clinical cases (Table S1). The majority of these strains (n=19) were part of the basic collection of Szmolka et al. (2018) describing the molecular epidemiology of *S*. Infantis between 2011 and 2013 in Hungary. Additionally, we included 12 broiler strains of *S*. Infantis isolated between 2016 and 2018, the latter being of caecal origin (see next section).

*E. coli* strains were isolated from the faeces, caecum and bone marrow of broiler chickens and day-old chicks (Table S1). Commensal strains from faeces (n=22) were isolated in 2013 from broiler chickens processed at three different slaughterhouses in North-Central Hungary. At slaughter 15 animals were randomly sampled at each slaughterhouse.

Caecal strains of *E. coli* (n=43) were isolated in 2018 from broiler chickens and from day-old chicks. Caecal samples from broiler chickens were provided by the Veterinary Diagnostic Directorate, National Food Chain Safety Office (NÉBIH), as part of the national *Salmonella* Monitoring Program (2018-2019). Here we tested 27 strains of *E. coli* isolated from the caecum of six broiler chickens, representing six different farms.

*E. coli* isolates from the caecum of day-old chicks (IntEC) were provided by colleagues from private veterinary services in Hungary (2018-2019). These strains were isolated from the caecum of nine chick carcasses, representing seven chicken farms (Table S1).

Extraintestinal pathogenic *E. coli* (ExPEC, n=25) strains were isolated from the bone marrow of day-old chicks that died from *E. coli* septicemia in two different periods. Old strains of ExPEC were isolated in Hungary between 1998-2000, and their virulence gene patterns were partly reported earlier (Tóth et al., 2012). Recent (2018) strains of ExPEC (2018) were isolated from the bone marrow of the same day-old chicks (n=9) from which the above mentioned IntEC strains also derived. Both, the caecal- and the bone marrow samples of day-old chicks were used to isolate multiple IntEC and ExPEC strains (Table S1), that were selected to represent antibiotic resistance phenotypes (see below). All *S*. Infantis and *E. coli* isolates were stored at −80°C in lysogeny broth (LB) (Becton Dickinson) containing 10% glycerol.

### 2.2. Isolation and identification of cohabitant strains of *S*. Infantis and *E. coli*

Caecal samples of broiler chickens (n=6) served as a common source for simultaneous isolation of *S*. Infantis and *E. coli* strains, that were designated here as cohabitant strains (Table S1). Each caecal sample was used to isolate multiple strains of *S*. Infantis and *E. coli*, to represent the intra-community resistance diversity and to predict potentially exchangeable genes and plasmid types between the two species.

Detection of *Salmonella* in the caecum of broilers was performed according to the international *Salmonella* standard ISO 6579:2006 with little modifications. Briefly, 10g caecal content was enriched in 100 ml LB broth for 18 hours at 37°C. After incubation, 10 ml RV (Rappaport-Vassiliadis) selective enrichment broth was inoculated with 100 µl of the overnight LB culture, and incubated for 24 hours at 42°C. Finally, 10 µl of the overnight RV culture was streaked onto XLD (Xylose Lysine Deoxycholate) agar plates (Becton Dickinson), and after 27 hours of incubation (37°C) ten individual *Salmonella*-like colonies were randomly picked to represent each sample. The molecular confirmation of *S*. Infantis was performed by serovar-specific PCR as described by Kardos et al. (2007).

Cohabitant strains of *E. coli* were isolated from the overnight LB cultures of the *Salmonella*-positive caecal samples. For this, 10 µl of the LB culture was streaked onto Chromocult® Coliform agar plates (Merck). After overnight incubation (18 hours, 37°C) ten individual *E. coli*-like colonies were picked randomly to represent each caecal sample. The molecular confirmation of all *E. coli* isolates was carried out by multiplex PCR on the basis of the simultaneous presence of the marker genes *lacZ* (beta-galactosidase) and *uidA* (beta-glucuronidase) (Bej et al., 1991).

### 2.3. Selection for multiresistance and determination of the pulsotype

Collections of *S*. Infantis and *E. coli* were assembled based on the antibiotic resistance of the strains to represent the diversity of phenotypes identified for different sample sources. For *S*. Infantis, the pulsotype was also considered by the selection of strains to represent clonal diversity.

The resistance phenotype was determined by disk diffusion against nine antibiotic compounds that were selected in order to detect plasmid-mediated phenotypes conferred by genes of the mobile resistomes. For this, the following antibiotic compounds were used: ampicillin (AMP_10_), cefotaxime (CTX_5_), chloramphenicol (CHL_30_), ciprofloxacin (CIP_5_), gentamicin (GEN_10_), meropenem (MEM_10_) nalidixic acid (NAL_30_), sulfonamide compounds (SUL_300_), tetracycline (TET_30_) and trimethoprim (TMP_5_). Antibiotic susceptibility testing was performed according to the guidelines and interpretation criteria of the European Committee on Antimicrobial Susceptibility Testing (EUCAST, 2018). *E. coli* ATCC 25922 was used as a reference strain. The intermediate category was interpreted as susceptible, while multiresistance was defined as simultaneous resistance against at least three antibiotic classes.

Pulsed-field gel electrophoresis (PFGE) of *S*. Infantis strains was carried out according to the CDC Pulse Net standardized *Salmonella* protocol using *S*. Braenderup H9812 as a molecular standard. *Xba*I restriction profiles were analysed by the Fingerprinting II Software (Bio-Rad Laboratories, Ventura, CA, USA). Cluster analysis was performed by the un-weighted pair-group method (UPGMA) with arithmetic means. Clonal distances were calculated on the basis of the Dice’s coefficient. A 1.0% position tolerance and 1.5% optimization setting were applied.

### 2.4. DNA extraction and whole-genome sequencing

Total genomic DNA was isolated using the MagAttract® HMW DNA kit (Qiagen, Hilden, Germany). To prepare 500 bp paired-end libraries of all isolates we used the Nextera XT DNA Library Preparation kit (Illumina, San Diego, CA, USA). Libraries were sequenced on the Illumina MiSeq sequencing platform using v2 sequencing chemistry. The quality of the raw sequencing data was then analyzed using FastQC v0.11.5 (Andrews, 2010). Raw reads were trimmed using Sickle v1.33 (https://github.com/najoshi/sickle). Genome assembly of the processed reads was carried out with SPAdes v3.10.1 (Bankevich et al., 2012).

### 2.5. *In silico* genome analysis

The *Salmonella* serovar identity was confirmed *in silico* on the basis of contig sequences by using the web-based application SeqSero v1.2 (Zhang et al., 2015). The *S*. Infantis-specific antigenic profile 7:r:1,5 (O:H1:H2) was thereby predicted for all the 56 *Salmonella* strains, perfectly corresponding to the Infantis-specific PCR. The *E. coli* serotype was determined by SerotypeFinder v.1.1 (Joensen et al., 2015) by using threshold values: identity 90%; minimum length 80%. The *E. coli* phylogroup was predicted by the *In Silico* Clermont Phylotyper (Waters et al., 2018) for the differentiation of the seven phylogroups (A, B1, B2, C, D, E, and F).

For molecular epidemiological analysis of *S*. Infantis and *E. coli* strains the Ridom SeqSphere+ software was used (Jünemann et al., 2013). Sequence types (STs) were determined by multilocus sequence typing (MLST) based on the polymorphism of the seven housekeeping genes according to the Warwick MLST scheme for *E. coli* (Wirth et al., 2006) and to the Achtman MLST scheme for *S*. Infantis (Achtman et al., 2012). Strains with unknown allelic profiles were submitted to Enterobase (Zhou et al., 2019) for the confirmation and identification of the new STs. The relatedness of *S*. Infantis and *E. coli* strains was further analysed by core genome (cg)MLST. The genome of the earliest sequenced *S*. Infantis strain 1326/28 (GenBank accession no. LN649235) was used as reference for the cgMLST of *S*. Infantis, while core genes of *E. coli* were analysed by blasting all genome sequences against the *E. coli* reference strain K-12 MG1655 (GenBank accession no. NC_000913).

Web-based tools ResFinder v.3.0 (Zankari et al., 2012) and PlasmidFinder v2.0 (Carattoli et al., 2014) were used for the *in silico* detection of acquired antibiotic resistance genes and for typing of plasmids on the basis of the replicon type. Selected thresholds for the prediction of resistance genes and replicon types were 90-95% sequence identity and 80% minimum length coverage. Cluster analysis based on the resistance genes was performed by using the PAST software v.4 (Hammer et al., 2001).

In order to reveal whether certain clones are associated with an increased prevalence of resistance genes the Multiple Antibiotic Resistance (MAR) index was calculated for all *E. coli* STs. The MAR index is the ratio between the total number of resistance genes, number of antibiotics tested and number of isolates (Krumperman, 1983). A higher MAR index indicates a greater abundance of resistance genes.

## 3. Results

### 3.1. Diversity of serotypes, STs and phylogroups of *E. coli* and of *S*. Infantis from broilers and humans

The SerotypeFinder predicted a large diversity of serotypes among the tested *E. coli* strains. Strains were assigned to 40 different O-types, out of which O8 and O9 were most frequently identified in eight and seven strains, respectively. The O-antigen-associated genes *wzx*/*wzy* were not typable (nt) for additional eight strains (Table S2). Similar to the O-serotype determinants, the H-type gene (*fliC*) was also variable, resulting in the identification of 31 H-types in the *E. coli* collection. The *in silico* phylogroup prediction resulted in the identification of predominant phylogroups A and B1 in 27% and 37% of the strains, respectively, most of them being isolated from the caecum. *E. coli* serotypes were compared to the phylogroup and the ST of the strains with regard to the source of isolation. In general, there was no strong correlation found between serotypes and phylogroups or sequence types of these *E. coli* strains. Serogroups O78, O88 and O115 showed a tendency for extraintestinal (bone marrow) isolates designated as ExPEC (Table S2).

In order to compare the diversity of the isolated *E. coli* and *S*. Infantis strains, the sequence types (STs) of 90 *E. coli* and of 56 S. Infantis strains were determined by MLST on the basis of whole genome sequences.

The *E. coli* strains were allocated to 49 STs (Fig. 1). The majority of them represented individual STs with one isolate each. Sequence types containing at least four strains were regarded here as large STs. Commensal and extraintestinal strains were both grouped in larger STs such as ST10, ST93, ST117 and ST162, while ST155 contained commensal (faecal, caecal) strains only. The new *E. coli* STs ST8702 and ST10088 representing caecal isolates belong to the clonal complexes CC10 and CC155, respectively (Table S1, Fig. 1).

**Figure 1.**
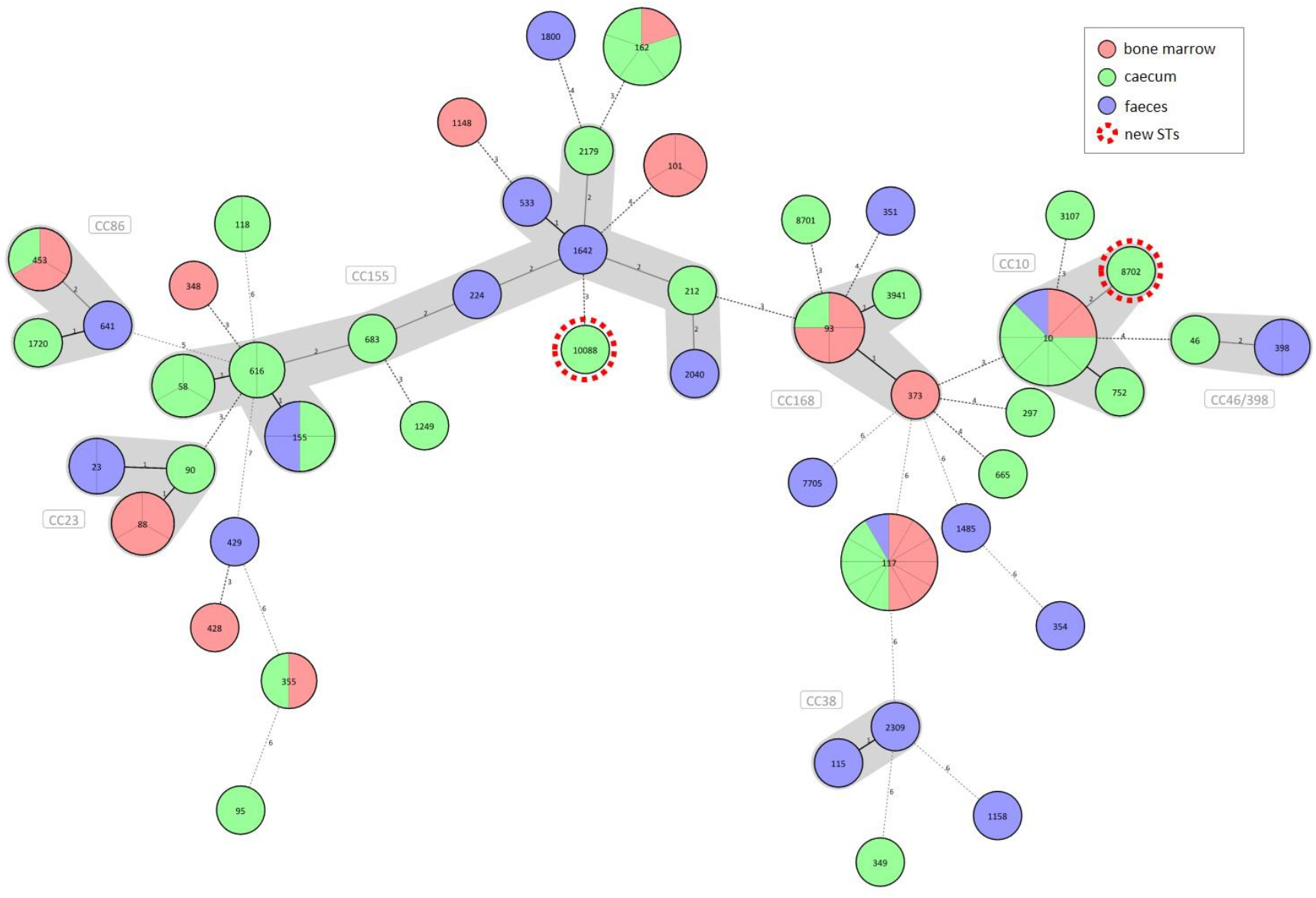
Diversity of commensal (caecal, faecal) and extraintestinal (bone marrow) *E. coli* strains from broilers according to the sample source. A Minimum Spanning Tree of 90 commensal and clinical *E. coli* strains was generated based on the polymorphism of seven housekeeping genes. Strains are separated by grey lines in nodes with multiple strains. Allelic patterns with missing values represented a category of its own and were designated as novel clones highlighted with red dotted circles. Distance lines change from black to dotted grey as the phylogenetic distance between the strains increases. Maximum distance in cluster was set to 2, making it therefore possible to group STs into clonal complexes (CCs). The thickness of the distance line is inversely proportional to the distance value.

In contrast to *E. coli*, most *S*. Infantis strains were assigned to ST32. Exceptions were two strains for which ST7081 (from human) and ST7082 (from broiler) were identified as new *Salmonella spp*. STs (Table S1).

### 3.2. Genomic diversity of *E. coli* and *S*. Infantis strains from broilers

To reveal genomic diversity of broiler strains of *E. coli*, cgMLST was carried out based on the polymorphism of 2398 genes of the core genome, by using the strain *E. coli* K-12 MG1655 as a reference. The core genome-tree showed that *E. coli* genomes are grouped into three main clusters according to their phylogenetic background (Fig. 2). Cluster 1 and Cluster 2 represent the dominant phylogroups A and B1 (27.8% and 37.8%), while most strains of Cluster 3 were assigned to phylogroup F (14.4%). Closely related strains from the large STs ST10, ST93 (Cluster 1) and ST155, ST162 (Cluster 2) showed a high level of genomic diversity on the basis of the core genome comparison. Commensal (intestinal) and extraintestinal (clinical) *E. coli* strains of ST117 displayed the most homogenous core genome sequences (Fig. 2). ST10 and ST93 belong to phylogroup A, ST117 to phylogroup F, while ST155 and ST162 belong to phylogroup B1 (Fig. 2).

**Figure 2.**
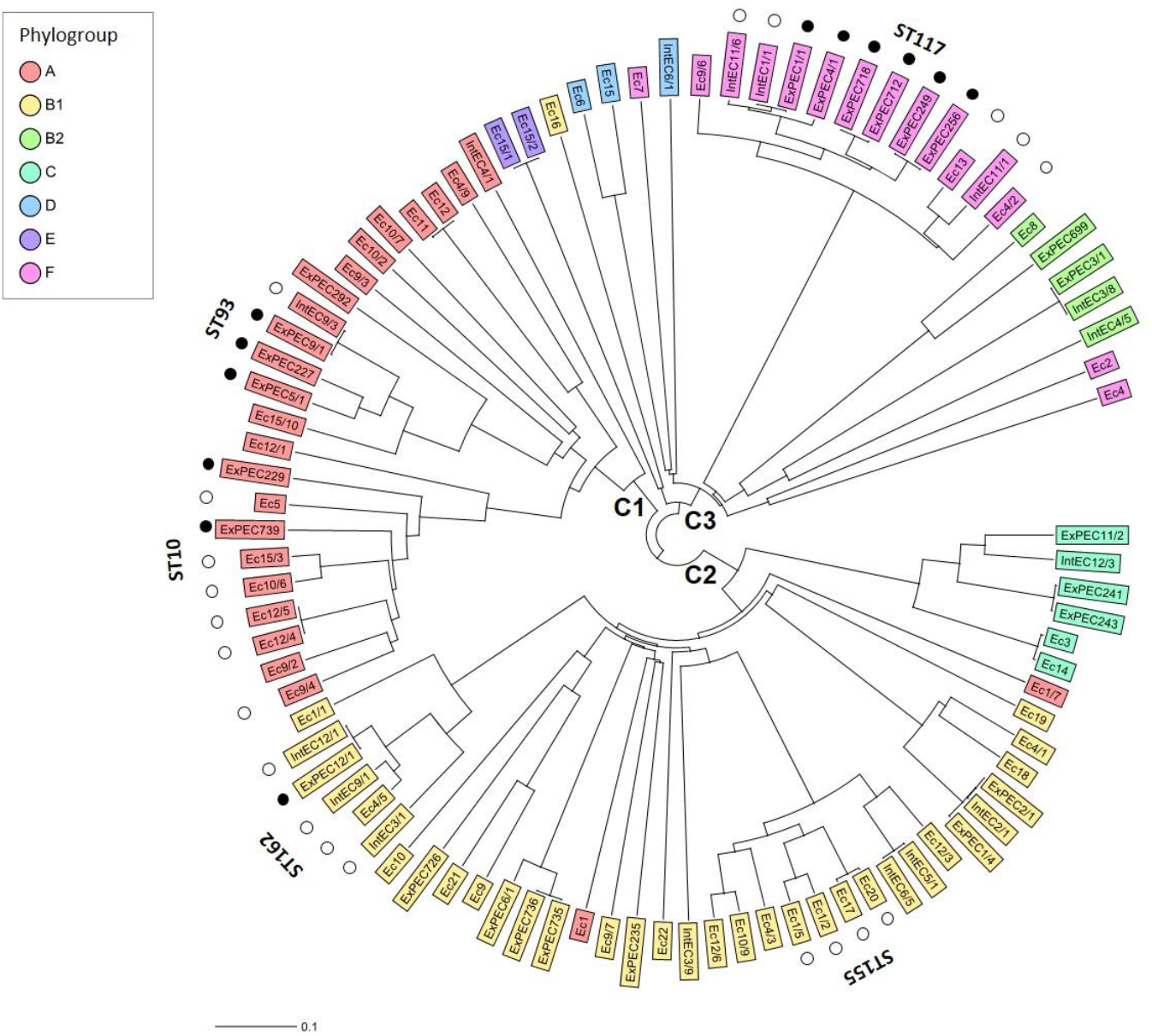
Core genome diversity and phylogroups of commensal (caecal, faecal) and extraintestinal (bone marrow) strains of *E. coli* from broilers. The Neighbour Joining Tree showing the genomic diversity of 90 *E. coli* strains was calculated based on the polymorphism of 2398 target genes of the core genome. Core genes were identified by blasting all genome sequences against the reference strain *E. coli* K-12 MG1655 (GenBank accession no. NC_000913). By distance calculation, gene columns with missing values were removed. Only the large STs of *E. coli* with at least four isolates are indicated. Black and white dots indicate extraintestinal and commensal (caecal, faecal) strains, respectively, representing larger STs. C1-3 describes Clusters 1-3.

For *S*. Infantis, the core genome analysis was performed to reveal the genomic diversity within the PFGE clones (Fig. 3), with special regard to the epidemic PFGE clone B2 (Table S1). The cgMLST was based on the sequence polymorphism of 3850 core genome targets by using the strain *S*. Infantis 1326/28 as a reference. Core genomes of 56 *S*. Infantis strains were analysed in comparison with the *S*. Infantis strains SI69/94 and SI54/04 representing the ancient, pansensitive strains of PFGE clone (pulsotype) A and of the emergent epidemic multiresistant PFGE clone B2, respectively.

**Figure 3.**
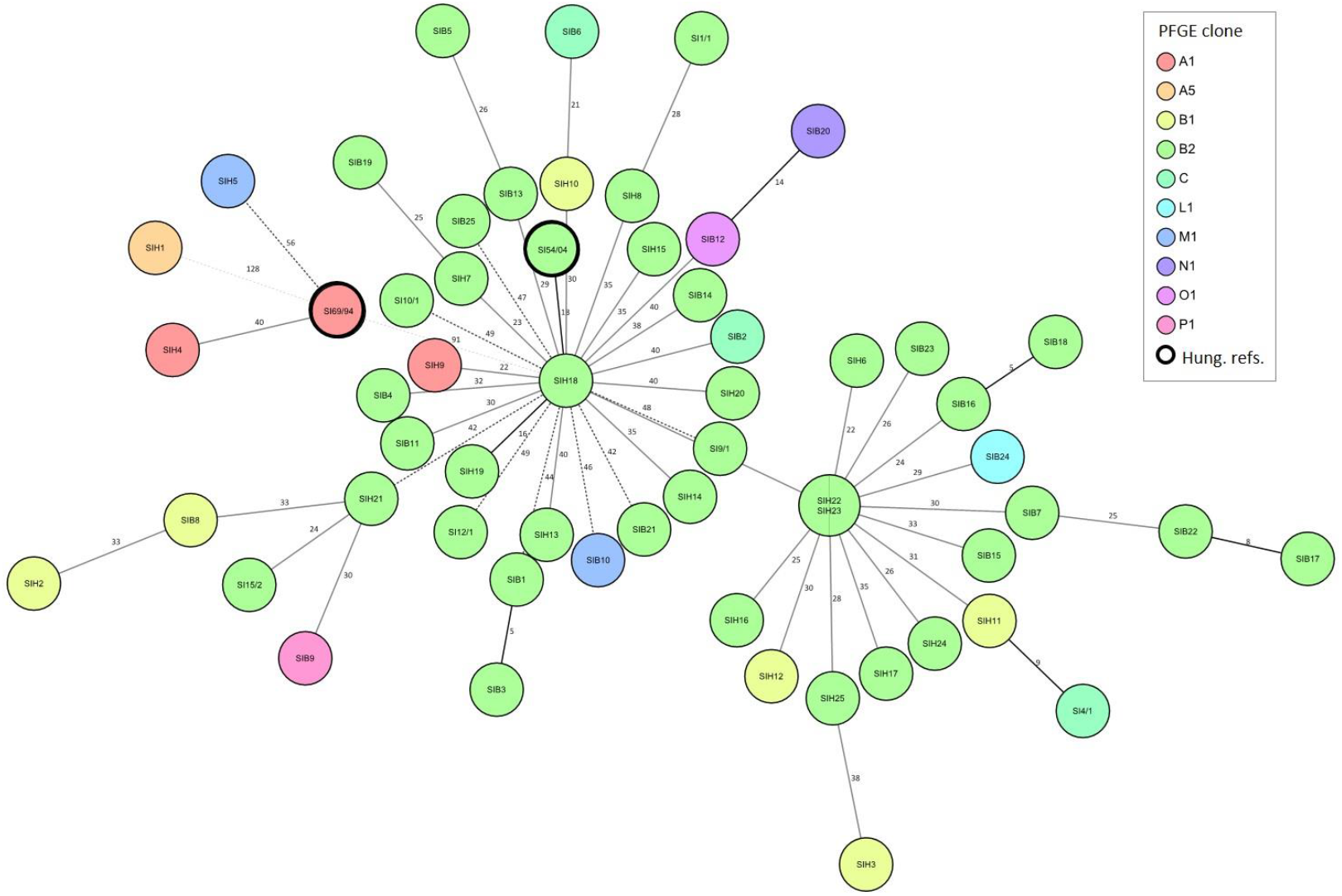
Relation between the genomic diversity and pulsotype in broiler and human strains of *S*. Infantis. Minimum Spanning Tree showing the core genome diversity of *S*. Infantis strains was generated based on the polymorphism of 3647 target genes of the core genome. By distance calculation, gene columns with missing values were removed. Distances are shown by the line style, distance numbers are also indicated. Distance lines change from black to dotted grey as the phylogenetic distance between the strains increases. Logarithmic scale was used for distance line length calculation. Strains are separated by grey lines in nodes with multiple strains. Besides the 56 *S*. Infantis strains studied here, the whole-genome sequences of *S*. Infantis strains SI69/94 (GenBank accession no. JRXB00000000) and SI54/04 (GenBank accession no. JRXC00000000) were also included, and blasted against the reference strain *S*. Infantis 1326/28 (GenBank accession no. LN649235). Both of these *S*. Infantis strains (framed by thick black line) were included as a Hungarian references for the ancient, pansensitive strains of the late 90’s (SI69/94) and for strains representing the emergent endemic MDR clone B2 (SI54/04) (Olasz et al., 2015).

The cgMLST analysis showed a high level of genomic diversity within the epidemic clone B2 of *S*. Infantis (Fig. 3). There were no relations between cgMLST and pulsotype. The *S*. Infantis strains grouped in two large clusters comprising strains from broilers and humans, and centred around the human strains SIH22/SIH23 or SIH18. This latter cluster contained the Hungarian MDR reference strain SI54/04. Core genomes of tested strains of *S*. Infantis strongly differed from that of the ancient reference strain SI69/94 (PFGE clone A). Exceptions were the human strains SIH1, SIH4 and SIH5 that clustered together with this ancient, pansensitive strain SI69/94, and one of them also represented the PFGE clone A.

### 3.3. The mobile resistome of *E. coli* and S. Infantis: differences and overlaps

To describe the diversity and distribution of acquired resistance genes and of plasmid types, the genome sequences of the tested *E. coli* and *S*. Infantis strains were analysed by the web-based programs ResFinder and PlasmidFinder. The *in silico* analysis showed that mobile resistomes of *E. coli* and *S*. Infantis strains could be compared along with a set of 17 resistance gene families (34 genes) and of 14 plasmid types (Table 1, Fig. 4).

**Table 1.**
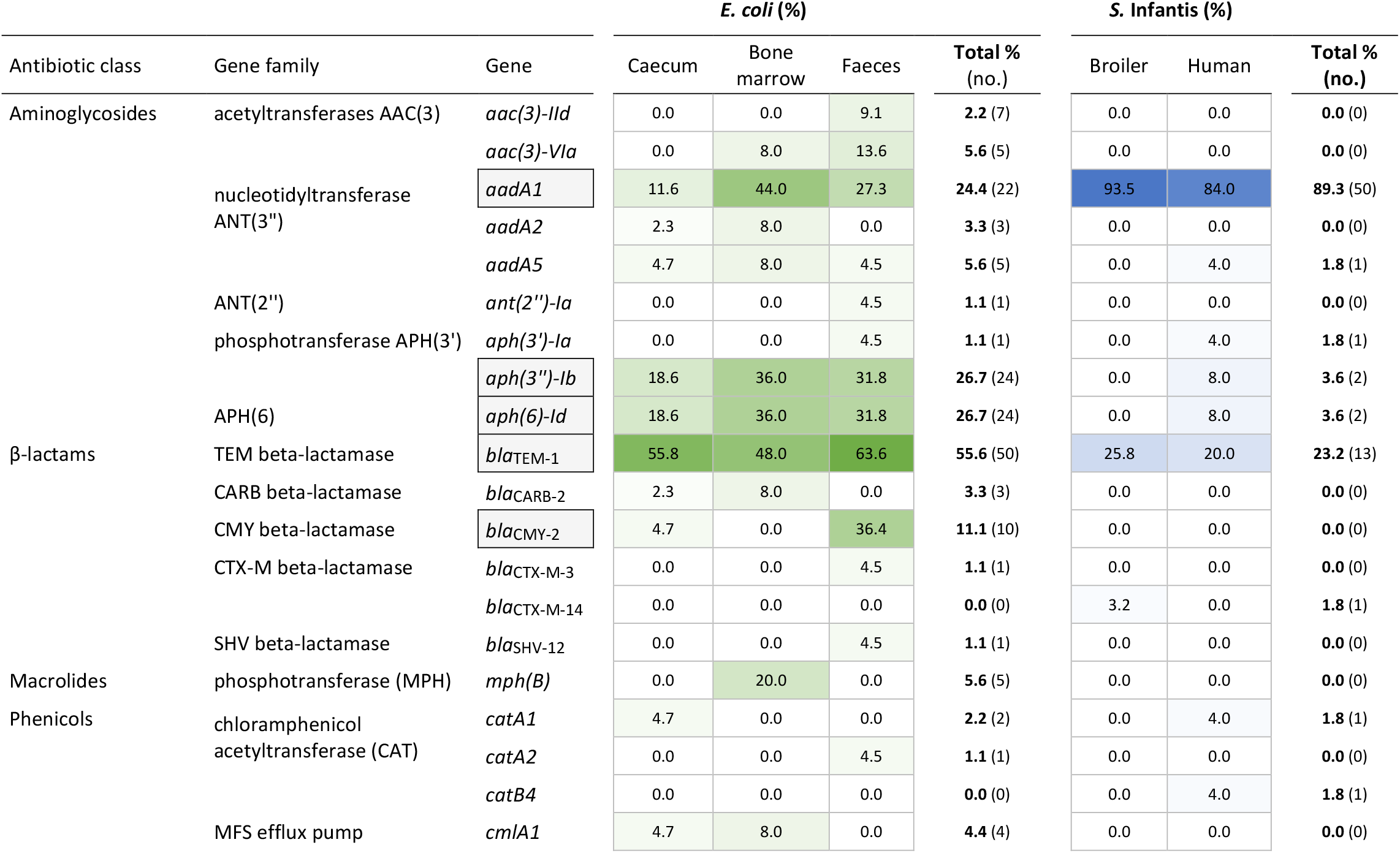
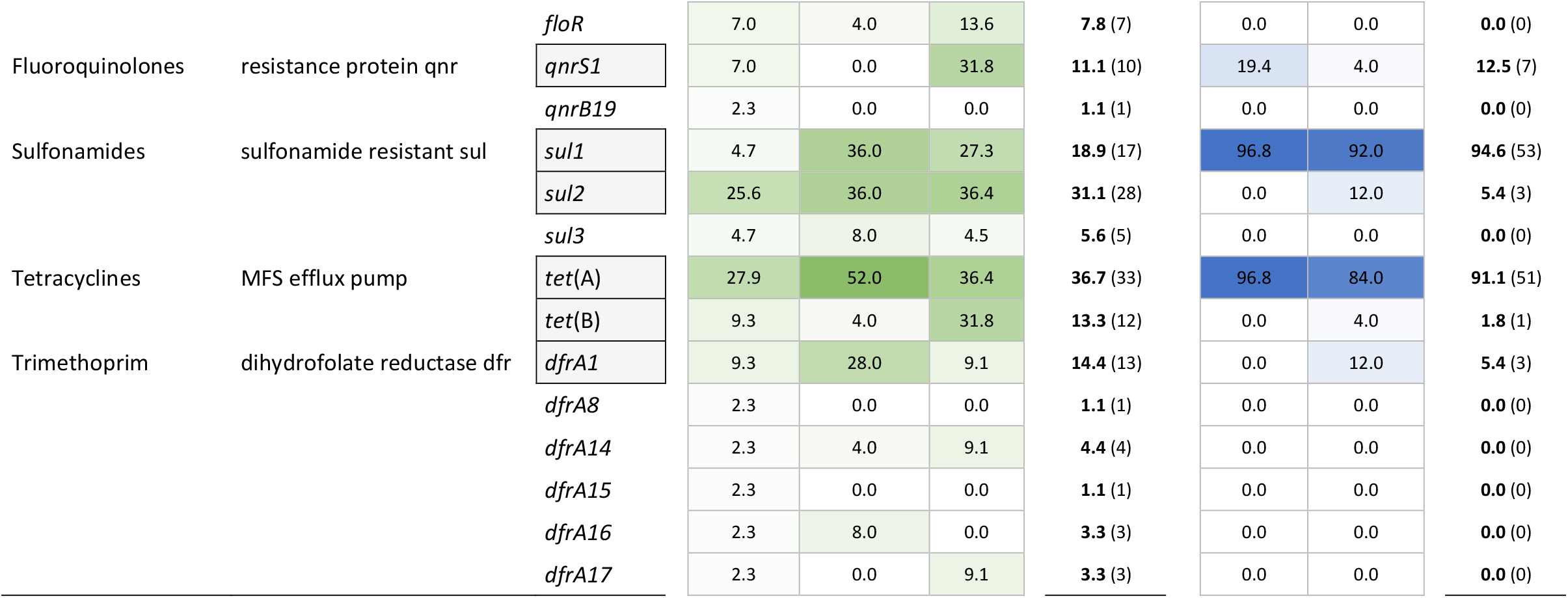
Diversity and distribution of resistance genes of the mobile resistomes in *S*. Infantis and *E. coli* strains isolated from broilers and humans. Grey background indicates genes present in >10% of the *E. coli* and *S*. Infantis strains. The intensity of green (*E. coli*) and blue (*S*. Infantis) colours increases with the frequency (%) of detection.

**Figure 4.**
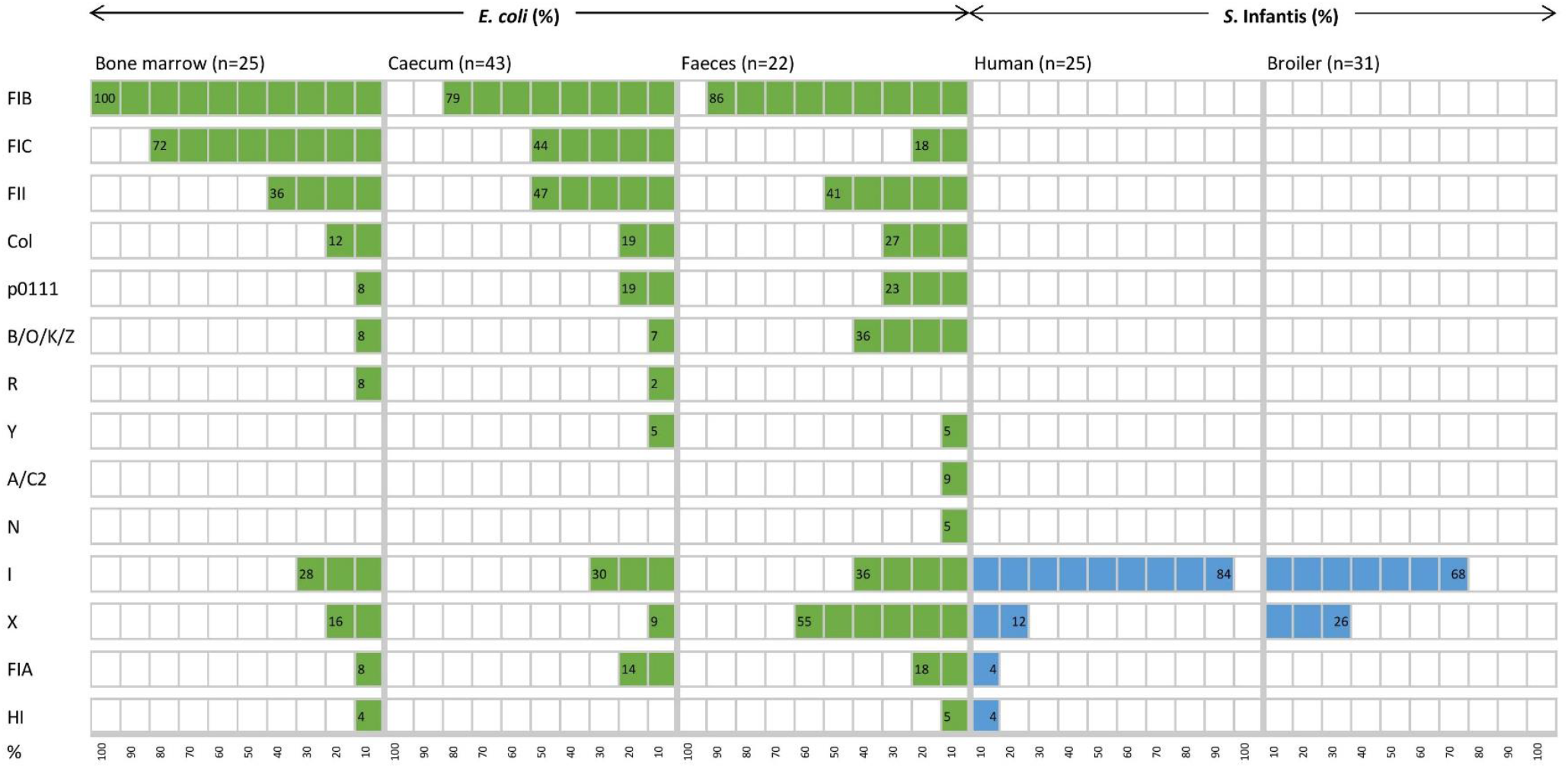
Diversity and distribution (%) of plasmid replicon types of *E. coli* and *S*. Infantis strains from different host sources.

The mobile resistome analysis of *E. coli* showed a large diversity of acquired resistance genes and plasmid types in the majority of *E. coli* strains. A total of 32 resistance genes were identified in the collection representing all the 17 gene families (Table 1). 11 of these genes were detected in more than 10% of the *E. coli* strains, conferring resistance to aminoglycosides, β-lactams, fluoroquinolone, sulfonamides, tetracyclines and trimethoprim. The distribution of resistance genes was not related to the sample type, but it seemed that *E. coli* strains isolated from faeces of slaughtered broilers and from bone marrow of day-old chicks showed a higher prevalence and diversity of resistance genes than those isolated from the caecum of day-old chicks (Table 1).

The ampicillin resistance gene *bla*_TEM-1_ and the tetracycline resistance gene *tet*(A) were most frequently detected, with a prevalence of 48.0-63.6% and 36.4-52.0% according to the source of isolation (Table 1). Extraintestinal strains (from bone marrow) demonstrated the highest frequency of genes *aadA1* (44.4%), *sul1* (36.0%) and *dfrA1* (28.0%) related to class 1 integrons. Besides, the macrolide phosphotransferase gene *mph*(B) was also detected exclusively in five extraintestinal *E. coli* strains. It seemed, however, that commensal (especially the faecal) *E. coli* strains rather carried genes related to emerging plasmids such as *qnrS1* (31.8%) and *bla*_CMY-2_ (36.4. The latter was identified only in *E. coli* strains. In comparison with *S*. Infantis, *E. coli* strains were also abundantly carrying genes coding for aminoglycoside-, sulphonamide- and tetracycline resistances. In this context, the *aph(3’’)-Ib, aph(6)-Id, sul2* and *tet*(B) genes should be mentioned as characterizing 13.3-26.7% of the *E. coli* strains, but being carried by only a few *S*. Infantis strains (Table 1).

In contrast, broiler and human strains of *S*. Infantis carried a more reduced set of resistance genes and plasmid types (Table 1, Fig. 4). A total of 15 resistance genes were identified in the collection representing ten gene families and five of these genes were detected in more than 10% of the *S*. Infantis strains, conferring resistance to aminoglycosides, β-lactams, fluoroquinolone, sulfonamides and tetracyclines (Table 1). The diversity of resistance genes was higher in human strains than in broiler strains. The tetracycline resistance gene *tet*(A) and the class 1 integron genes *aadA1* and *sul1* were predominantly identified (89.3-91.1%) in *S*. Infantis strains regardless of their host. These genes were also frequently identified in *E. coli* but at a considerably lower prevalence (Table 1). Additional resistance genes detected in *E. coli* as well as in *S*. Infantis were the ampicillin- and fluoroquinolone resistance genes *bla*_TEM-1_ (23.2%) and *qnrS1* (12.5%), respectively. The prevalence of *bla*_TEM-1_ was much higher in *E. coli* than in *S*. Infantis, while *qnrS1* gene was detected more frequently in broiler strains than in human strains of *S*. Infantis. The emerging CTX-M type resistance genes were also present in both species. According to this, the *bla*_CTX-M-14_ gene was identified in one broiler faecal strain of *S*. Infantis, while *bla*_CTX-M-3_ was detected in one faecal *E. coli* strain (Table 1).

Consistent with the high diversity of resistance genes, the coexistence of multiple plasmids was predicted for the majority of *E. coli* strains. Most plasmids belonged to the F replicon type, and it seemed that plasmid diversity was higher in commensal strains than in extraintestinal ones (Fig. 4). Because at least two different plasmid types were detected in almost all strains, no correlation could be established between certain resistance genes and plasmid types, except for one strain which carried the *bla*_TEM-1_ gene on an IncI plasmid (data not shown).

In contrast to *E. coli*, most *S*. Infantis strains were identified as monoplasmidic, and the coexistence of two replicon types was predicted for only two strains from humans. The MDR plasmid pSI54/04 of *S*. Infantis dominant in Hungary (Szmolka et al., 2018) was identified with the replicon type I, similar to the CTX-M-14 plasmid carried by a broiler *S*. Infantis isolate. Despite of the above mentioned remarkable differences regarding the plasmid diversity of the tested *E. coli* and *S*. Infantis strains, plasmids with replicon types I and X were most commonly detected in both species (Fig. 4).

We also wanted to find out if increased resistance was associated with certain *E. coli* STs. As a result, we determined that altogether 10 STs (including up to three strains each) proved to comprise potential carriers of multiresistence (MAR index >0.3) (Fig. S1).

### 3.4. Relations between resistance genes and plasmids of *E. coli* and *S*. Infantis and comparison of their cohabitant strains

To obtain deeper insight into resistance relationships in *E. coli* and *S*. Infantis, the strains were grouped according to the pattern of acquired resistance genes. This cluster analysis showed that the composition of the *E. coli* resistomes was very heterogenous, even with the association of up to 10 resistance genes in some of the strains (Fig. 5). According to the resistance phenotype, MDR strains of *E. coli* were characterized by several combinations of associated resistance genes. Based on these constellations, *E. coli* strains were grouped into 3 well-separable clusters: i) cluster CARB characterized by the coexistence of genes *bla*_CARB-2_ -*bla*_TEM-1_, ii) cluster TEM/TET with the associated genotype of *bla*_TEM-1_ – *tet*(A) – *qnrS*, and iii) cluster APH, that could be regarded as a “ super-MDR clade” as it is characterised by multiple combinations of associated genes around the *aph(3’’)-Ib* and *aph(6)-Id* genes. Genes related to class 1 integrons were commonly detected in all of these three clusters. (Fig. 5).

**Figure 5.**
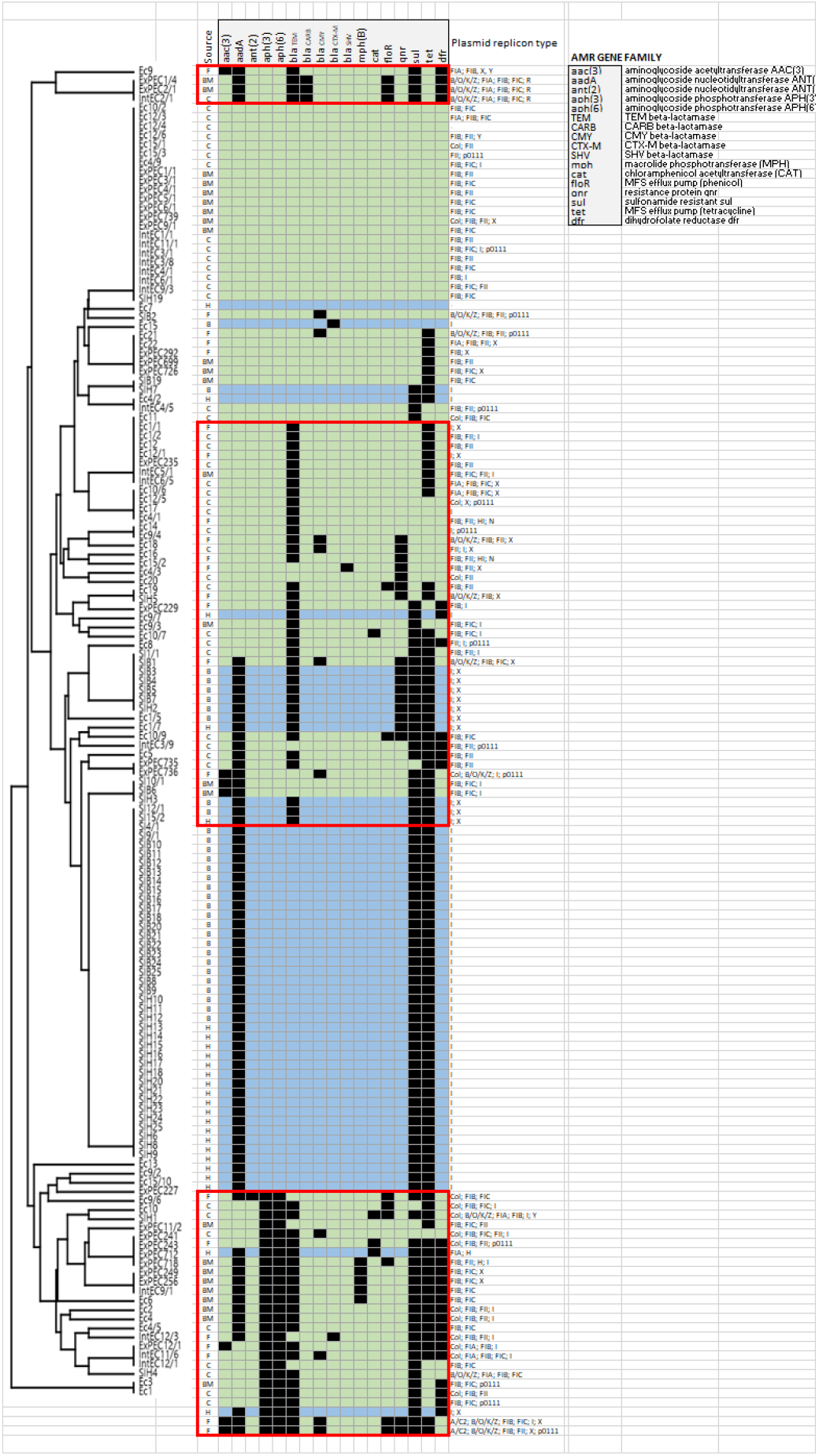
Mobile resistome tree of *S*. Infantis isolated from broilers and humans and of *E. coli* from broilers. *E. coli* strains are coloured in light green, while light blue represents *S*. Infantis strains. Abbreviations for the source of isolation are: F (faeces); BM (bone marrow); C (caecum); B (broiler) and H (human). The identified *E. coli* clusters are framed by red lines.

In contrast, the composition of the of *S*. Infantis resistomes was quite homogenous, with the association of max. five genes in the majority of strains. Most *S*. Infantis isolates from broilers and humans grouped into a large cluster designated here as cluster TET, characterized by the MDR genotype *tet*(A)-*aadA1*-*sul1*. Some strains isolated from broilers were grouped into the clade TEM/TET identified for *E. coli*. Accordingly, these strains carried the *bla*_TEM-1_ – *qnrS1* genes in addition to the *tet*(A)-*aadA1*-*sul1* genes. The simultaneous presence of six to eight resistance genes was detected in only two human *S*. Infantis isolates, both of them were grouped in the above mentioned “ super-MDR clade” identified for *E. coli* (Fig. 5).

For the prediction of potentially exchangeable resistance genes/plasmids the comparison of the mobile resistomes of cohabitant *E. coli* and *S*. Infantis strains was performed. Cohabitant strains of *E. coli* and *S*. Infantis were isolated from the same samples, more exactly from the caecum of six broiler chickens representing six different farms (Fig. 6). From each caecal sample it was possible to isolate one MDR strain of *S*. Infantis together with at least four cohabitant *E. coli* strains with different resistance phenotypes. The comparative resistome analysis of the cohabitant strains showed that the class 1 integron genes *aadA-sul1* and the resistance plasmid genes *tet*(A), *bla*_TEM-1_ and *qnrS1* can be potentially exchangeable between *E. coli* and *S*. Infantis. Furthermore, certain plasmids, especially those with replicon type I could be involved in the interaction between these two species (Fig. 6).

**Figure 6.**
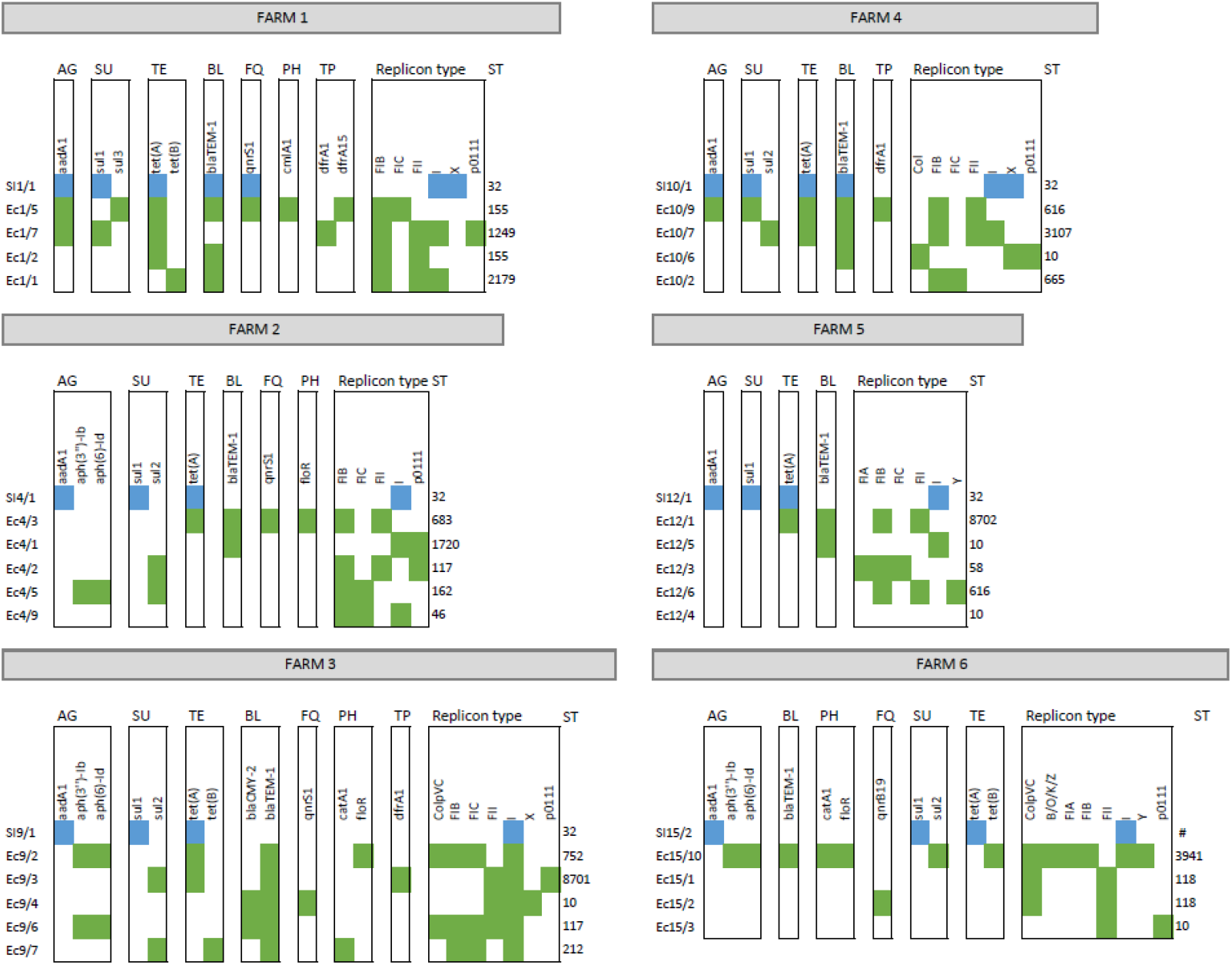
Antibiotic resistance genes as potentially exchangeable between cohabitant strains of *E. coli* and *S*. Infantis from caecal samples of broilers. *E. coli* strains are coloured in green, while blue represents *S*. Infantis strains. Abbreviations for antibiotic resistances are: AG (aminoglycosides); SU (sulfonamides); TE (tetracyclines); BL (beta-lactams); FQ (fluoroquinolones); PH (phenicols) and TP (trimethoprim).

## 4. Discussion

In the last decade the multiresistance of *S*. Infantis has grown into a global problem of the broiler industry. MDR strains of *S*. Infantis and *E. coli* strains are frequently co-isolated with MDR *E. coli* from broilers, which increases the food safety risk of chicken meat and contributes to decreasing its consumer value and market potential.

The simultaneous presence of both bacterial species may allow an active intraspecies and intergeneric interplay of mobile resistance elements between *S*. Infantis and *E. coli* in broilers. Here we provided a genome-based comparison of mobile resistomes and genomic diversity of *S*. Infantis and *E. coli* isolates in order to gather new information about relations between these two cohabitant MDR enterobacteria of food safety importance. The core genome-based analysis identified a high genomic heterogeneity of broiler *E. coli* strains.

This high-resolution analysis allowed the first insight into the relatedness of *S*. Infantis isolates of PFGE clone B2, which is epidemic in Hungary among broilers and humans. At the same time, we were able to provide an overview on emerging lineages and mobile resistance determinants characterizing populations of *E. coli* and *S*. Infantis in Hungarian broiler flocks. By the comparative analysis of mobile resistomes in cohabitant strains, the first snapshot was taken on a possible intergeneric interaction and resistance gene exchange between *E. coli* and *S*. Infantis in the caecum of chickens.

Genomic diversity of *E. coli* was not related to the source of isolation. Strains from both, caecum and bone marrow, displayed a high diversity in STs and serotypes. None of the *E. coli* STs represented MDR high-risk lineages, such as ST131 known to cause extraintestinal infection of humans, but also colonizing several animal hosts (Mathers et al., 2015; Yamaji et al., 2018b). ST10, ST93, ST117 and ST162 were identified here as larger closely related groups, comprising both intestinal and extraintestinal strains, while ST155 contained intestinal strains only. ST10 is known as one of the most widespread MDR lineages of *E. coli* from animals (Liu et al., 2018), and can also be isolated from urine of patients suffering from urinary tract infection (Yamaji et al., 2018a). ST93 was described among others to include ESBL-/and *mcr-1* expressing *E. coli* isolates from foods of animal origin (Zhang et al., 2019). Similarly, *mcr-1* positive *E. coli* strains of ST162 were isolated from food in China (Zhang et al., 2019) and from pigs in Mexico (Rueda Furlan et al., 2019). Furthermore, ST117 and ST155 have also been recently reported to include many human pathogenic and MDR *E. coli* strains with zoonotic potential (Alonso et al, 2017; Yang et al., 2017). ST117 was also described as the primary causative agent of cellulitis in poultry in Denmark (Poulsen et al., 2018). Here, we identified two new MDR clones, i.e. ST8702 and ST10088, of the clonal complexes (CCs) CC10 and CC155, respectively. The above mentioned ST117 and ST155 could not be considered as reservoirs for genetic determinants of high priority resistance.

Regarding the O-type diversity, we most frequently identified O8, O9 and O78 in *E. coli* strains from broilers. From among these, the O78 group is widely recognized for ExPEC in chickens and man (Schouler et al., 2012; Tóth et al., 2012) and as such, may constitute a zoonotic risk (Moulin-Schouleur et al., 2007; Huja et al., 2015). Moreover, it seems that avian *E. coli* O78:H4/ST117 detected here in the bone marrow and in caecal contents could represent an epidemic lineage, as several reports indicate that O78:H4 is an epidemiologically significant serotype of avian ExPEC representing the ST117 (Ronco et al., 2017; Poulsen et al., 2018). Although, *E. coli* strains of serogroup O8 also seem to be implicated in avian colibacillosis (Gross et al., 1991), much less is known about extraintestinal pathogenicity of this group of *E. coli* isolates in the avian host (Schouler et al., 2012). This is in good agreement with our finding that only one of the eight MDR *E. coli* O8 strains was derived from the bone marrow. There seem to be no data available on avian *E. coli* O9 strains, which were all caecal (commensal) isolates in our study. The presence of these three O serogroups (especially of O78) will justify an in depth analysis of virulence genes of these *E. coli* isolates, which should be the subject of further genomic analysis in the near future.

It could also be stated that strains from the large STs such as ST10, ST93 (Cluster 1) and ST155, ST162 (Cluster 2) displayed a high level of genomic diversity on the basis of the core genome comparison, but *E. coli* ST117 strains were the most homogenous in their genome content (Cluster 3). Finally, it should also be mentioned that ExPEC strains were relatively most frequently grouped to ST117 in Cluster 3 (six of 14 such strains: 42.8%) in line with previous data on avian pathogenic *E. coli* of ST117 (Alonso et al, 2017; Yang et al., 2017; Poulsen et al, 2018). This contrasts with the lower proportional occurrence of ExPEC among phylogroup A strains of Cluster 1 (six of 23 such strains: 26.1%) and of ExPEC among phylogroup B1 strains of Cluster 2 (eight of 33 such strains: 24.2%).

In contrast to the heterogeneity of *E. coli* described above, strains of *S*. Infantis have shown a very homogenous clonal structure by almost all being assigned to ST32, which is considered the predominant ST of this serovar worldwide. However, it appears that *S*. Infantis is genetically “ open” for a clonal diversification, as indicated by the emergence of MDR STs such as the ST2283, which has recently spread in the broiler population in Germany (Garcia-Soto et al., 2020). Our data also contribute to the global clonal diversity of *S*. Infantis through the identification of two new MDR STs ST7081 and ST7082 for a human and a broiler strain, respectively.

The predominant prevalence of the MDR genotype of *tet*(A)-*aadA1*-*sul1* indicates the constant circulation of the dominant PFGE clone B2 and its MDR plasmid in Hungary since its emergence two decades ago (Nógrády et al., 2007; Szmolka et al., 2018). By using the high discriminatory cgMLST genotyping tool, we revealed for the first time the internal genomic structure of the epidemic PFGE clone B2, showing two main clusters each with a high genomic plasticity and potential for clonal diversification. This finding is in line with other reports, stating that the *S*. Infantis population is heterogeneous at the genomic level, with a more homogeneous repertoire of MDR plasmids, and with a potential to reflect geographical differences (Gymoese et al., 2019; Alba et al., 2020; Nagy et al., 2020).

Our results from comparative mobile resistome analysis describes avian *E. coli* strains as genetically diverse with a high prevalence and diversity of plasmid types and of mobile resistance genes, while *S*. Infantis strains proved to be much poorer in antibiotic resistance determinants and in different plasmid types. Such a discrepancy between these two bacterial populations is reflecting the genetic differences between these two species that may be explained by their divergent evolution about 100-150 million years ago (Ochman and Wilson, 1987). Besides, *E. coli* is a genetically extremely flexible long-term intestinal colonizer and member of the normal intestinal microbiota, perfectly adapted to changing conditions in the intestinal tract, while *S*. Infantis is a rather transient invader in the gut, having to compete with the resident microbiota of chicks.

We identified plasmid replicon types in the *S*. Infantis and *E. coli* isolates to get an overview of the interspecies distribution of plasmid types without the need to assign individual resistance genes to plasmid types. From all the plasmid types detected in our study, IncF plasmids represented the most striking difference between *E. coli* and *S*. Infantis. IncF plasmids are common broad-host-range plasmids of enteric bacteria associated with virulence in pathogenic *E. coli* (Johnson and Nolan, 2009). They are also well-known as MDR plasmids. Moreover, the emergence of some dominant *E. coli* STs has been driven by ESBL plasmids of the IncF family (Villa et al., 2010; Carattoli, 2013; Dunn et al., 2019). We detected this replicon family almost exclusively in *E. coli* strains with the exception of one human *S*. Infantis isolate, suggesting that the prevalence of this plasmid type is greatly determined by the bacterial host species. The IncF plasmid family has rarely been reported in *Salmonella*, e.g. the *spvB* virulence plasmid of *S*. Typhimurium belongs to the IncF plasmid family (Oluwadare et al., 2020). IncI plasmids, however, were predominantly identified in *S*. Infantis, but this plasmid family was also frequently detected in *E coli* strains.

For both species the mobile resistance genes *tet*(A), *aadA1* and *sul1* were identified with an overall high prevalence, conferring resistance to classical antibiotic classes such as tetracyclines, aminoglycosides and sulfonamides, while the *bla*_TEM-1_ gene encoding ampicillin resistance was detected mostly in *E. coli* regardless of the sample source. These findings on the most frequently identified resistance genes fully support the latest EU report on the prevalence of antibiotic resistance phenotypes in poultry (EFSA, 2020). The abundance of these genes and of the corresponding MDR genotypes is not surprising, as a number of compounds belonging to these antibiotic classes are approved and largely used in some EU countries for disease prevention in broilers (Roth et al., 2019). Thus, the coexistence of genes *bla*_TEM-1,_ *tet*(A), *aadA1* and *sul1* is not something particularly noteworthy for *E. coli*, and the combination of these associations is increasingly considered as a common feature of both commensal and pathogenic strains (Szmolka et al., 2013; Vila et al., 2016). But for *S*. Infantis, the *tet*(A)-*aadA1-sul1* genotype indicated the emergence of a new epidemic MDR clone (pulsotype B2) of *S*. Infantis, that was first reported in Hungary (Nógrády et al., 2007; Nógrády et al., 2012) and subsequently in other countries (Franco et al., 2015; Hindermann et al., 2017).

The epidemiological success and persistence of the clone B2 still seems to be unbroken, as we predominantly identified the MDR genotype *tet*(A)-*aadA1*-*sul1* among broiler and human strains of *S*. Infantis. This is in line with our previous report, in which we traced the molecular epidemiology of Hungarian strains of *S*. Infantis between 2011 and 2013 (Szmolka et al., 2018). The coexistence of these mobile resistance marker genes and of the specific virulence genes identified previously (Szmolka et al., 2018) indicated the indisputable presence of the plasmid pSI54/04 in most of the *S*. Infantis strains. Plasmid pSI54/04 was first defined as an IncP plasmid by the PCR-based replicon typing system developed by Carattoli et al. (2005), however, the WGS-based pMLST (Carattoli et al., 2014) identified this plasmid as belonging to the IncI1 family.

Plasmid pSI54/04 is regarded as a pESI-like plasmid (<u>p</u>lasmid for <u>e</u>merging *<u>S</u>*. Infantis), because of showing high sequence similarity with the resistance-virulence megaplasmid pESI endemic in *S*. Infantis in Israel (Aviv et al., 2014; Szmolka et al; 2018). The real incompatibility group of some pESI-like plasmids is not clearly definable, because the replication origin of these plasmids is not conventional. It results from the substitution of the IncI1 *oriV* by an IncP-1alpha *oriV* (Aviv et al., 2014; Dionisi et al., 2016). This could be one reason for the controversial findings described above regarding incompatibility of pSI54/04, basically confirming findings of Bogomazova et al. (2020) in relation to comparative analysis of a selected set of pESI-like plasmid-bearing strains on the basis of whole-genome sequencing. It is conceivable that this non-conventional replicon type may be an advantage for pESI-like plasmids, in facilitating selection and global spread of certain successful MDR clones of *S*. Infantis such as the clone B2, which is epidemic in Hungary.

pESI-like plasmids of *S*. Infantis gain further importance, because of their ability to coexist with certain resistance plasmids and/or to incorporate multiple resistance genes including those conferring resistance to extended-spectrum β-lactamases (ESBLs) (Bogomazova et al., 2020).

As an example for plasmid coexistence, we describe the association between the tetracycline resistance plasmid pSI54/04 and a small *bla*_TEM-1_-*qnrS1* plasmid of IncX incompatibility group responsible for β-lactam- and fluoroquinolone resistance in a small group of broiler strains. Kehrenberg et al. (2006) described for the first time a conjugative *bla*_TEM-1_-*qnrS1* plasmid pINF5 in a chicken isolate of *S*. Infantis, but the emergence of new alleles such as *bla*_TEM-20_, *bla*_TEM-52_, *bla*_TEM-70_, *bla*_TEM-148_ and *bla*_TEM-198_ seems to diminish the scientific importance of the wild-type TEM-1 plasmids for this serovar (Cloeckaert et al., 2007; Shahada et al., 2010; Chuma et al., 2013). Very recently, the coexistence of a pESI-like megaplasmid with an IncX plasmid carrying the the *mcr-1* gene coding for colistin resistance has been reported in *S*. Infantis isolated from broilers (Carfora et al., 2018). Moreover, the ESBL gene *bla*_CTX-M-1_ was identified on a pESI-like plasmid of the IncP incompatibility group. Indeed, some successful MDR clones of *S*. Infantis in Europe and the United States carry pESI-like plasmids conferring ESBL resistance by genes *bla*_CTX-M-1_ and *bla*_CTX-M-65_ (Franco et al., 2015; Tate et al., 2017; Hindermann et al., 2017; Alba et al., 2020). In contrast, the plasmid pSI54/04 is constantly and predominantly present in the Hungarian broiler and human *S*. Infantis population, but does not carry ESBL genes.

In comparison to the overall high-level resistance to the above mentioned “ classical” antibiotics, resistances to high-priority antibiotic classes such as fluoroquinolones, third-generation cephalosporins, carbapenems, polymixins and macrolides were detected at a lower level or were not detected at all. The plasmid-mediated fluoroquinolone resistance gene *qnrS1* characterized our broiler intestinal strains of *E. coli* and *S*. Infantis relatively frequently, but it was absent from extraintestinal *E. coli* strains. In harmony with these findings, the EU member states also reported a high prevalence of fluoroquinolone resistance in commensal (indicator) *E. coli* from broilers (EFSA, 2020). The *E. coli* resistance to third-generation cephalosporins was conferred here by the ESBL genes *bla*_CMY-2_ and *bla*_CTX-M-3_ detected in a small number of strains with origin from faeces and caecum. In Europe, resistance to cephalosporins is also reported at a low incidence (EFSA, 2020), but in some countries outside Europe the intestine of the broiler chickens is regarded as a reservoir of ESBL-producing *E. coli* (Li et al., 2016; Vinueza-Burgos et al., 2019). For *S*. Infantis, we report the first ESBL-producing human strain in Hungary that carries the cefotaxime resistance gene *bla*_CTX-M-14_ on an IncI1 type plasmid. This allele is rarely identified in broiler strains of *S*. Infantis (Kameyama et al., 2012; Bogomazova et al., 2020), and it is more likely embedded in conjugative plasmids of *S*. Enteritidis of human origin (Romero et al., 2004; Izumiya et al., 2005; Bado et al., 2012). The mobile resistance gene *mph(B)* encoding a macrolide efflux pump was identified in five ExPEC strains that were grouped in the so-called “ super-MDR clade” characterized by multiple combinations of associated resistance genes and mechanisms. Besides, certain ExPEC strains are effective in causing severe diseases in chickens and posing a serious concern to food safety and to human health (Gao et al., 2018).

The WGS-based comparison of antibiotic resistance genes and plasmid types revealed a relatively narrow interface between the mobile resistomes of *E. coli* and of *S*. Infantis. According to this, we found that the resistance genes *tet*(A)-*aadA1*-*sul1* specific to the S. Infantis plasmid pSI54/04 were also frequently identified in *E. coli* strains, but the association of genes *bla*_TEM-1_ and *qnrS1* were also commonly carried by members of both species. By the comparative mobile resistome analysis of the cohabitant strains, we were able to confirm the presence of these genes and of plasmid types IncI and IncX in strains of *E. coli* and *S*. Infantis isolated from the same caecum of chicken. These findings suggest that the potential for an interspecies transmission exists for certain resistance genes or even for whole plasmids such as the pSI54/04 or the *bla*_TEM-1_ plasmid. Indeed, it was demonstrated that the pESI plasmid endemic for *S*. Infantis in Israel can be transferred *in vivo* to *E. coli* strains as members of the normal gut microflora in mouse (Aviv et al., 2016), but so far there is a general lack of knowledge regarding the transferability of pESI-like plasmids of *S*. Infantis to *E. coli* in broilers.

It seems that the comparative characterization of cohabitant strains is a promising approach, that would improve our understanding of the epidemiology of plasmids with zoonotic significance. However, our results so far can be considered as preliminary and conjugation experiments along this line are representing our future task.

## 5. Conclusions

This is the first report on the comparative genomic analysis of contemporary strains of *S*. Infantis and of *E. coli* from broilers. The present analysis confirms the continuous persistence and reveals the high genomic diversity of multiresistant *E. coli* and *S*. Infantis endemic in broiler flocks in Hungary. Our findings on the diversity of the mobile resistomes indicate that commensal *E. coli* can be regarded as a potential reservoir of resistance genes for *Salmonella*, but so far only a few plasmid types and genes of mobile resistomes of *E. coli* and *S*. Infantis could be considered as potentially exchangeable. Among these, certain IncI and IncX plasmids could play the most important role in resistance gene/plasmid exchange and the evolution of multiresistance between *E. coli* and *S*. Infantis. Future active molecular monitoring of MDR *E. coli* and cohabitant *S*. Infantis strains will certainly shed more light on the microevolution of emerging clones and plasmids of MDR *S*. Infantis in broilers.

## 6. Author contributions

AS and UD conceived the project and wrote the manuscript. HW and AS performed the bioinformatic and resistome analyses, respectively. All authors reviewed the manuscript.

## 8. Acknowledgments

The authors thank Prof. Béla Nagy for his essential help during this project and for the critical review of the manuscript. Thanks are also due to Dr. Szilárd Jánosi and Judit Szolyák for providing the chicken caecal samples for the isolation of cohabitant strains of *S*. Infantis and *E. coli*. We also thank Dr. Judit Pászti for providing the PFGE analysis for *S*. Infantis strains isolated in 2016 and 2018. Further thanks are due to Dr. Csaba Szalay and Dr. Béla Markos for providing *E. coli* cultures from the caecum and the bone marrow of chicks. Haleluya Wami was supported by a PhD scholarship of Pharma-Zentrale GmbH (Herdecke). We thank Karin Tegelkamp for the excellent performance of the genome sequencing of the *S*. Infantis and *E. coli* isolates.

## References

Acar, S., Bulut, E., Stasiewicz, M. J., and Soyer, Y. (2019). Genome analysis of antimicrobial resistance, virulence, and plasmid presence in Turkish Salmonella serovar Infantis isolates. Int J Food Microbiol 307, 108275. doi:10.1016/j.ijfoodmicro.2019.108275.

Achtman, M., Wain, J., Weill, F.-X., Nair, S., Zhou, Z., Sangal, V., et al. (2012). Multilocus Sequence Typing as a Replacement for Serotyping in Salmonella enterica. PLoS Pathog 8, e1002776. doi:10.1371/journal.ppat.1002776.

Ahmed, A. M., Shimabukuro, H., and Shimamoto, T. (2009). Isolation and Molecular Characterization of Multidrug-Resistant Strains of Escherichia coli and Salmonella from Retail Chicken Meat in Japan. J Food Sci 74, M405–M410. doi:10.1111/j.1750-3841.2009.01291.x.

Alba, P., Leekitcharoenphon, P., Carfora, V., Amoruso, R., Cordaro, G., Matteo, P. D., et al. (2020). Molecular epidemiology of Salmonella Infantis in Europe: insights into the success of the bacterial host and its parasitic pESI-like megaplasmid. Microb Genom. doi:10.1099/mgen.0.000365.

Alonso, C. A., Michael, G. B., Li, J., Somalo, S., Simón, C., Wang, Y., et al. (2017). Analysis of blaSHV-12-carrying Escherichia coli clones and plasmids from human, animal and food sources. J Antimicrob Chemoth 72, 1589–1596. doi:10.1093/jac/dkx024.

Andrews, S. (2010). FastQC: a quality control tool for high throughput sequence data. Available online at: http://www.bioinformatics.babraham.ac.uk/projects/fastqc

Aviv, G., Rahav, G., and Gal-Mor, O. (2016). Horizontal Transfer of the Salmonella enterica Serovar Infantis Resistance and Virulence Plasmid pESI to the Gut Microbiota of Warm-Blooded Hosts. mBio 7, e01395–16. doi:10.1128/mbio.01395-16.

Aviv, G., Tsyba, K., Steck, N., Salmon-Divon, M., Cornelius, A., Rahav, G., et al. (2014). A unique megaplasmid contributes to stress tolerance and pathogenicity of an emergent Salmonella enterica serovar Infantis strain. Environ Microbiol 16, 977–94. doi:10.1111/1462-2920.12351.

Bado, I., García-Fulgueiras, V., Cordeiro, N. F., Betancor, L., Caiata, L., Seija, V., et al. (2012). First Human Isolate of Salmonella enterica Serotype Enteritidis Harboring blaCTX-M-14 in South America. Antimicrob Agents Chemother 56, 2132–2134. doi:10.1128/aac.05530-11.

Bankevich, A., Nurk, S., Antipov, D., Gurevich, A. A., Dvorkin, M., Kulikov, A. S., et al. (2012). SPAdes: A new genome assembly algorithm and its applications to single-cell sequencing. Journal of Computational Biology, 19(5), 455–477. doi:10.1089/cmb.2012.0021

Bej, A. K., McCarty, S. C., and Atlas, R. M. (1991). Detection of coliform bacteria and Escherichia coli by multiplex polymerase chain reaction: comparison with defined substrate and plating methods for water quality monitoring. Applied and environmental microbiology 57, 2429–32.

Blake, D. P., Hillman, K., Fenlon, D. R., and Low, J. C. (2003). Transfer of antibiotic resistance between commensal and pathogenic members of the Enterobacteriaceae under ileal conditions. J Appl Microbiol 95, 428–436. doi:10.1046/j.1365-2672.2003.01988.x.

Bogomazova, A. N., Gordeeva, V. D., Krylova, E. V., Soltynskaya, I. V., Davydova, E. E., Ivanova, O. E., et al. (2019). Mega-plasmid found worldwide confers multiple antimicrobial resistance in Salmonella Infantis of broiler origin in Russia. Int J Food Microbiol 319, 108497. doi:10.1016/j.ijfoodmicro.2019.108497.

Carattoli, A., Zankari, E., García-Fernández, A., Larsen, M. V., Lund, O., Villa, L., et al. (2014). In silico detection and typing of plasmids using PlasmidFinder and plasmid multilocus sequence typing. Antimicrob Agents Chemother 58, 3895–903. doi:10.1128/AAC.02412-14.

Carattoli, A. (2013). Plasmids and the spread of resistance. Int J Med Microbiol 303, 298–304. doi:10.1016/j.ijmm.2013.02.001.

Carattoli, A., Bertini, A., Villa, L., Falbo, V., Hopkins, K. L., and Threlfall, E. (2005). Identification of plasmids by PCR-based replicon typing. J Microbiol Methods 63, 219–28. doi:10.1016/j.mimet.2005.03.018.

Carfora, V., Alba, P., Leekitcharoenphon, P., Ballarò, D., Cordaro, G., Matteo, P. D., et al. (2018). Colistin Resistance Mediated by mcr-1 in ESBL-Producing, Multidrug Resistant Salmonella Infantis in Broiler Chicken Industry, Italy (2016–2017). Front Microbiol 9, 1880. doi:10.3389/fmicb.2018.01880.

Chuma, T., Miyasako, D., Dahshan, H., Takayama, T., Nakamoto, Y., Shahada, F., et al. (2013). Chronological Change of Resistance to β-Lactams in Salmonella enterica serovar Infantis Isolated from Broilers in Japan. Front Microbiol 4, 113. doi:10.3389/fmicb.2013.00113.

Cloeckaert, A., Praud, K., Doublet, B., Bertini, A., Carattoli, A., Butaye, P., et al. (2007). Dissemination of an Extended-Spectrum-β-Lactamase blaTEM-52 Gene-Carrying IncI1 Plasmid in Various Salmonella enterica Serovars Isolated from Poultry and Humans in Belgium and France between 2001 and 2005. Antimicrob Agents Chemother 51, 1872–1875. doi:10.1128/AAC.01514-06.

Dionisi, A. M., Owczarek, S., Benedetti, I., Luzzi, I., and García-Fernández, A. (2016). Extended-spectrum β-lactamase-producing Salmonella enterica serovar Infantis from humans in Italy. Int J Antimicrob Agents 48, 345–6. doi:10.1016/j.ijantimicag.2016.06.025.

Dunn, S. J., Connor, C., and McNally, A. (2019). The evolution and transmission of multi-drug resistant Escherichia coli and Klebsiella pneumoniae: the complexity of clones and plasmids. Curr Opin Microbiol 51, 51–56. doi:10.1016/j.mib.2019.06.004.

EUCAST. The European Committee on Antimicrobial Susceptibility Testing. Breakpoint tables for interpretation of MICs and zone diameters. Version 8.0. (2018) http://www.eucast.org.

European Food Safety Authority [EFSA] and European Centre for Disease Prevention [ECDC] (2020). The European Union Summary Report on Antimicrobial Resistance in zoonotic and indicator bacteria from humans, animals and food in 2017/2018. EFSA J 18, e06007. doi:10.2903/j.efsa.2020.6007.

Franco, A., Leekitcharoenphon, P., Feltrin, F., Alba, P., Cordaro, G., Iurescia, M., et al. (2015). Emergence of a Clonal Lineage of Multidrug-Resistant ESBL-Producing Salmonella Infantis Transmitted from Broilers and Broiler Meat to Humans in Italy between 2011 and 2014. PLoS One 10, e0144802. doi:10.1371/journal.pone.0144802.

Furlan, J. P. R., Gallo, I. F. L., Campos, A. C. L. P. de, Passaglia, J., Falcão, J. P., Navarro, A., et al. (2019). Molecular characterization of multidrug-resistant Shiga toxin-producing Escherichia coli harboring antimicrobial resistance genes obtained from a farmhouse. Pathog Glob Health 113, 1–7. doi:10.1080/20477724.2019.1693712.

Gal-Mor, O., Valinsky, L., Weinberger, M., Guy, S., Jaffe, J., Schorr, Y., et al. (2010). Multidrug-Resistant Salmonella enterica Serovar Infantis, Israel. Emerg Infect Dis 16, 1754–1757. doi:10.3201/eid1611.100100.

Gao, J., Duan, X., Li, X., Cao, H., Wang, Y., and Zheng, S. J. (2018). Emerging of a highly pathogenic and multi-drug resistant strain of Escherichia coli causing an outbreak of colibacillosis in chickens. Infect Genetics Evol 65, 392–398. doi:10.1016/j.meegid.2018.08.026.

García-Soto, S., Abdel-Glil, M. Y., Tomaso, H., Linde, J., and Methner, U. (2020). Emergence of Multidrug-Resistant Salmonella enterica Subspecies enterica Serovar Infantis of Multilocus Sequence Type 2283 in German Broiler Farms. Front Microbiol 11, 1741. doi:10.3389/fmicb.2020.01741.

Gross W.B. (1991). Colibacillosis, p.138–144. In: Calnek B.W., Barnes H.J., Beard C.W., Reid W.M. & Yoder J.H.W. (Eds), Disease of Poultry. 9th ed. Iowa State University Press, Ames.

Gyles, C. L. (2008). Antimicrobial resistance in selected bacteria from poultry. Anim Health Res Rev. 9, 149–158. doi:10.1017/s1466252308001552.

Gymoese, P., Kiil, K., Torpdahl, M., Østerlund, M. T., Sørensen, G., Olsen, J. E., et al. (2019). WGS based study of the population structure of Salmonella enterica serovar Infantis. BMC Genomics 20, 870. doi:10.1186/s12864-019-6260-6.

Hammer, Ø., Harper, D.A.T., Ryan, P.D. (2001). PAST: Paleontological statistics software package for education and data analysis. Palaeontol Electron 4(1): 9pp.

Hindermann, D., Gopinath, G., Chase, H., Negrete, F., Althaus, D., Zurfluh, K., et al. (2017). Salmonella enterica serovar Infantis from Food and Human Infections, Switzerland, 2010–2015:

Poultry-Related Multidrug Resistant Clones and an Emerging ESBL Producing Clonal Lineage. Front Microbiol 08, 1322. doi:10.3389/fmicb.2017.01322.

Huja, S., Oren, Y., Trost, E., Brzuszkiewicz, E., Biran, D., Blom, J., et al. (2015). Genomic Avenue to Avian Colisepticemia. Mbio 6, e01681–14. doi:10.1128/mbio.01681-14.

Izumiya, H., Mori, K., Higashide, M., Tamura, K., Takai, N., Hirose, K., et al. (2005). Identification of CTX-M-14 β-Lactamase in a Salmonella enterica Serovar Enteritidis Isolate from Japan. Antimicrob Agents Chemother 49, 2568–2570. doi:10.1128/aac.49.6.2568-2570.2005.

Joensen, K. G., Tetzschner, A. M. M., Iguchi, A., Aarestrup, F. M., and Scheutz, F. (2015). Rapid and Easy In Silico Serotyping of Escherichia coli Isolates by Use of Whole-Genome Sequencing Data. J Clin Microbiol 53, 2410–26. doi:10.1128/jcm.00008-15.

Johnson, T. J., and Nolan, L. K. (2009). Pathogenomics of the Virulence Plasmids of Escherichia coli. Microbiol Mol Biol Rev 73, 750–774. doi:10.1128/mmbr.00015-09.

Jünemann, S., Sedlazeck, F. J., Prior, K., Albersmeier, A., John, U., Kalinowski, J., et al. (2013). Updating benchtop sequencing performance comparison. Nat Biotechnol 31, 294–6. doi:10.1038/nbt.2522.

Kameyama, M., Chuma, T., Yokoi, T., Yata, J., Tominaga, K., Miyasako, D., et al. (2012). Emergence of Salmonella enterica Serovar Infantis Harboring IncI1 Plasmid with blaCTX-M-14 in a Broiler Farm in Japan. J Vet Med Sci 74, 1213–1216. doi:10.1292/jvms.11-0488.

Kardos, G., Farkas, T., Antal, M., Nógrády, N., and Kiss, I. (2007). Novel PCR assay for identification of Salmonella enterica serovar Infantis. Letters in Applied Microbiology 45, 421– 425. doi:10.1111/j.1472-765X.2007.02220.x.

Kehrenberg, C., Friederichs, S., Jong, A. de, Michael, G. B., and Schwarz, S. (2006). Identification of the plasmid-borne quinolone resistance gene qnrS in Salmonella enterica serovar Infantis. J Antimicrob Chemother 58, 18–22. doi:10.1093/jac/dkl213.

Krumperman, P. H. (1983). Multiple antibiotic resistance indexing of Escherichia coli to identify high-risk sources of fecal contamination of foods. Appl Environ Microbiol 46, 165–170.

Li, S., Zhao, M., Liu, J., Zhou, Y., and Miao, Z. (2016). Prevalence and Antibiotic Resistance Profiles of Extended-Spectrum β-Lactamase–Producing Escherichia coli Isolated from Healthy Broilers in Shandong Province, China. J Food Protect 79, 1169–1173. doi:10.4315/0362-028x.jfp-16-025.

Liu, X., Liu, H., Wang, L., Peng, Q., Li, Y., Zhou, H., et al. (2018). Molecular Characterization of Extended-Spectrum β-Lactamase-Producing Multidrug Resistant Escherichia coli From Swine in Northwest China. Front Microbiol 9, 1756. doi:10.3389/fmicb.2018.01756.

Mathers, A. J., Peirano, G., and Pitout, J. D. D. (2015). Chapter Four Escherichia coli ST131: The Quintessential Example of an International Multiresistant High-Risk Clone. Adv Appl Microbiol 90, 109–154. doi:10.1016/bs.aambs.2014.09.002.

Mathew, A.G., Jackson, F., and Saxton, A.M. (2002). Effects of antibiotic regimens on resistance of Escherichia coli and Salmonella serovar Typhimurium in swine. J Swine Health Prod 10, 7–13.

Moulin-Schouleur, M., Répérant, M., Laurent, S., Brée, A., Mignon-Grasteau, S., Germon, P., et al. (2007). Extraintestinal Pathogenic Escherichia coli Strains of Avian and Human Origin: Link between Phylogenetic Relationships and Common Virulence Patterns. J Clin Microbiol 45, 3366– 3376. doi:10.1128/jcm.00037-07.

Nagy, T., Szmolka, A., Wilk, T., Kiss, J., Szabó, M., Pászti, J., et al. (2020). Comparative Genome Analysis of Hungarian and Global Strains of Salmonella Infantis. Front Microbiol 11, 539. doi:10.3389/fmicb.2020.00539.

Nógrády, N., Király, M., Davies, R., and Nagy, B. (2012). Multidrug resistant clones of Salmonella Infantis of broiler origin in Europe. Int J Food Microbiol 157, 108–112. doi:10.1016/j.ijfoodmicro.2012.04.007.

Nógrády, N., Tóth, Á., Kostyák, Á., Pászti, J., and Nagy, B. (2007). Emergence of multidrug-resistant clones of Salmonella Infantis in broiler chickens and humans in Hungary. J Antimicrob Chemoth 60, 645–648. doi:10.1093/jac/dkm249.

Ochman, H., and Wilson, A. C. (1987). Evolution in bacteria: Evidence for a universal substitution rate in cellular genomes. J Mol Evol 26, 377–377. doi:10.1007/bf02101157.

Oladeinde, A., Cook, K., Lakin, S. M., Woyda, R., Abdo, Z., Looft, T., et al. (2019). Horizontal Gene Transfer and Acquired Antibiotic Resistance in Salmonella enterica Serovar Heidelberg following In Vitro Incubation in Broiler Ceca. Appl Environ Microb 85. doi:10.1128/aem.01903-19.

Olasz, F., Nagy, T., Szabó, M., Kiss, J., Szmolka, A., Barta, E., et al. (2015). Genome Sequences of Three Salmonella enterica subsp. enterica Serovar Infantis Strains from Healthy Broiler Chicks in Hungary and in the United Kingdom. Genome Announc 3, e01468–14. doi:10.1128/genomea.01468-14.

Oluwadare, M., Lee, M. D., Grim, C. J., Lipp, E. K., Cheng, Y., and Maurer, J. J. (2020). The Role of the Salmonella spvB IncF Plasmid and Its Resident Entry Exclusion Gene traS on Plasmid Exclusion. Front Microbiol 11, 949. doi:10.3389/fmicb.2020.00949.

Poppe, C., Martin, L. C., Gyles, C. L., Reid-Smith, R., Boerlin, P., McEwen, S. A., et al. (2005). Acquisition of Resistance to Extended-Spectrum Cephalosporins by Salmonella enterica subsp. enterica Serovar Newport and Escherichia coli in the Turkey Poult Intestinal Tract. Appl Environ Microb 71, 1184–1192. doi:10.1128/aem.71.3.1184-1192.2005.

Poulsen, L. L., Bisgaard, M., Jørgensen, S. L., Dideriksen, T., Pedersen, J. R., and Christensen, H. (2018). Characterization of Escherichia coli causing cellulitis in broilers. Vet Microbiol 225, 72– 78. doi:10.1016/j.vetmic.2018.09.011.

Romero, L., López, L., Martínez-Martínez, L., Guerra, B., Hernández, J. R., and Pascual, A. (2004). Characterization of the first CTX-M-14-producing Salmonella enterica serotype Enteritidis isolate. J Antimicrob Chemoth 53, 1113–1114. doi:10.1093/jac/dkh246.

Ronco, T., Stegger, M., Olsen, R. H., Sekse, C., Nordstoga, A. B., Pohjanvirta, T., et al. (2017). Spread of avian pathogenic Escherichia coli ST117 O78:H4 in Nordic broiler production. BMC Genomics 18, 13. doi:10.1186/s12864-016-3415-6.

Roth, N., Käsbohrer, A., Mayrhofer, S., Zitz, U., Hofacre, C., and Domig, K. J. (2019). The application of antibiotics in broiler production and the resulting antibiotic resistance in Escherichia coli: A global overview. Poultry Sci 98, 1791–1804. doi:10.3382/ps/pey539.

Schouler, C., Schaeffer, B., Brée, A., Mora, A., Dahbi, G., Biet, F., et al. (2012). Diagnostic Strategy for Identifying Avian Pathogenic Escherichia coli Based on Four Patterns of Virulence Genes. J Clin Microbiol 50, 1673–1678. doi:10.1128/jcm.05057-11.

Shahada, F., Chuma, T., Dahshan, H., Akiba, M., Sueyoshi, M., and Okamoto, K. (2010). Detection and characterization of extended-spectrum beta-lactamase (TEM-52)-producing Salmonella serotype Infantis from broilers in Japan. Foodborne Pathog Dis 7, 515–21. doi:10.1089/fpd.2009.0454.

Shahada, F., Chuma, T., Tobata, T., Okamoto, K., Sueyoshi, M., and Takase, K. (2006). Molecular epidemiology of antimicrobial resistance among Salmonella enterica serovar Infantis from poultry in Kagoshima, Japan. Int J Antimicrob Agents 28, 302–307. doi:10.1016/j.ijantimicag.2006.07.003.

Szmolka, A., and Nagy, B. (2013). Multidrug resistant commensal Escherichia coli in animals and its impact for public health. Front Microbiol 4, 258. doi:10.3389/fmicb.2013.00258.

Szmolka, A., Szabó, M., Kiss, J., Pászti, J., Adrián, E., Olasz, F., et al. (2018). Molecular epidemiology of the endemic multiresistance plasmid pSI54/04 of Salmonella Infantis in broiler and human population in Hungary. Food Microbiol 71, 25–31. doi:10.1016/j.fm.2017.03.011.

Tate, H., Folster, J. P., Hsu, C.-H. H., Chen, J., Hoffmann, M., Li, C., et al. (2017). Comparative Analysis of Extended-Spectrum-β-Lactamase CTX-M-65-Producing Salmonella enterica Serovar Infantis Isolates from Humans, Food Animals, and Retail Chickens in the United States. Antimicrob Agents Chemother 61, e0488–17. doi:10.1128/aac.00488-17.

Tóth, I., Dobrindt, U., Koscsó, B., Kósa, A., Herpay, M., and Nagy, B. (2012). Genetic and phylogenetic analysis of avian extraintestinal and intestinal Escherichia coli. Acta Microbiol Imm H 59, 393–409. doi:10.1556/amicr.59.2012.3.10.

Varga, C., Guerin, M. T., Brash, M. L., Slavic, D., Boerlin, P., and Susta, L. (2019). Antimicrobial resistance in fecal Escherichia coli and Salmonella enterica isolates: a two-year prospective study of small poultry flocks in Ontario, Canada. BMC Vet Res 15, 464. doi:10.1186/s12917-019-2187-z.

Vila, J., Sáez-López, E., Johnson, J., Römling, U., Dobrindt, U., Cantón, R., et al. (2016). Escherichia coli: an old friend with new tidings. FEMS Microbiol Rev 40, 437–463. doi:10.1093/femsre/fuw005.

Villa, L., García-Fernández, A., Fortini, D., and Carattoli, A. (2010). Replicon sequence typing of IncF plasmids carrying virulence and resistance determinants. J Antimicrob Chemoth 65, 2518– 2529. doi:10.1093/jac/dkq347.

Vinueza-Burgos, C., Ortega-Paredes, D., Narváez, C., Zutter, L. D., and Zurita, J. (2019). Characterization of cefotaxime resistant Escherichia coli isolated from broiler farms in Ecuador. PLos One 14, e0207567. doi:10.1371/journal.pone.0207567.

Waters, N., Brennan, F., Holmes, A., Abram, F., and Pritchard, L. (2018). Easily phylotyping E. coli via the EzClermont web app and command-line tool. bioRxiv, 317610. doi:10.1101/317610.

Wirth, T., Falush, D., Lan, R., Colles, F., Mensa, P., Wieler, L. H., et al. (2006). Sex and virulence in Escherichia coli: an evolutionary perspective. Mol Microbiol 60, 1136–1151. doi:10.1111/j.1365-2958.2006.05172.x.

Yamaji, R., Friedman, C. R., Rubin, J., Suh, J., Thys, E., McDermott, P., et al. (2018a). A Population-Based Surveillance Study of Shared Genotypes of Escherichia coli Isolates from Retail Meat and Suspected Cases of Urinary Tract Infections. Msphere 3, e00179–18. doi:10.1128/msphere.00179-18.

Yamaji, R., Rubin, J., Thys, E., Friedman, C. R., and Riley, L. W. (2018b). Persistent Pandemic Lineages of Uropathogenic Escherichia coli in a College Community from 1999 to 2017. J Clin Microbiol 56, e01834–17. doi:10.1128/jcm.01834-17.

Yang, Y.-Q., Li, Y.-X., Song, T., Yang, Y.-X., Jiang, W., Zhang, A.-Y., et al. (2017). Colistin Resistance Gene mcr-1 and Its Variant in Escherichia coli Isolates from Chickens in China. Antimicrob Agents Ch 61, e01204–16. doi:10.1128/aac.01204-16.

Zankari, E., Hasman, H., Cosentino, S., Vestergaard, M., Rasmussen, S., Lund, O., et al. (2012). Identification of acquired antimicrobial resistance genes. J Antimicrob Chemoth 67, 2640–2644. doi:10.1093/jac/dks261.

Zhang, S., Yin, Y., Jones, M. B., Zhang, Z., Kaiser, B. L., Dinsmore, B. A., et al. (2015). Salmonella Serotype Determination Utilizing High-Throughput Genome Sequencing Data. J Clin Microbiol 53, 1685–1692. doi:10.1128/JCM.00323-15.

Zhang, P., Wang, J., Wang, X., Bai, X., Ma, J., Dang, R., et al. (2019). Characterization of Five Escherichia coli Isolates Co-expressing ESBL and MCR-1 Resistance Mechanisms From Different Origins in China. Front Microbiol 10, 1994. doi:10.3389/fmicb.2019.01994.

Zhou, Z., Alikhan, N.-F., Mohamed, K., Fan, Y., Group, the A. S., Achtman, M., et al. (2019). The EnteroBase user’s guide, with case studies on Salmonella transmissions, Yersinia pestis phylogeny and Escherichia core genomic diversity. Genome Res 30, 138–152. doi:10.1101/gr.251678.119.

