## Supplemental Figure 1 for "Comparative genomics of emerging lineages and mobile resistomes of contemporary broiler strains of *Salmonella* Infantis and *E. coli*"

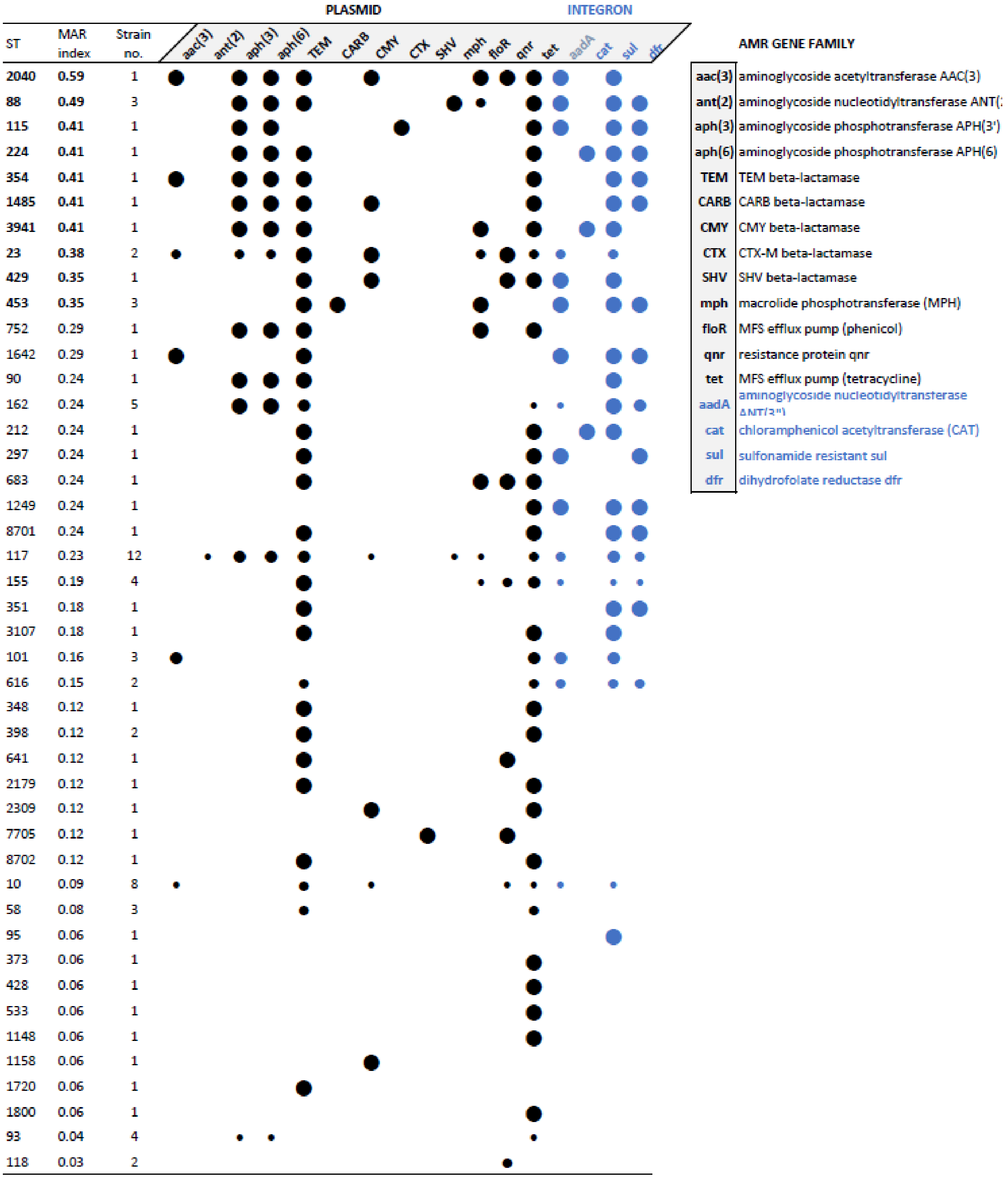


**Figure S1. Relation between the sequence type (ST) and abundance of acquired resistance genes.** In bold are indicated STs that are potentially associated with increased abundance of acquired resistance genes (MAR index >0.3). Circle sizes are proportional to the prevalence (%) of the given gene for each ST. "Empty" STs (ST46, 349, 355, 665) are not presented.
