## Supplemental Table 1 for "Comparative genomics of emerging lineages and mobile resistomes of contemporary broiler strains of *Salmonella* Infantis and *E. coli*"

**Table S1**. **Sequenced strains of broiler and human *S*. Infantis (SI) ■ and of broiler *E. coli* (Ec) ■ with origin from different sample sources.**
Black dots indicate cohabitant strains of SI and Ec isolated from six caecal samples representing one farm each. Cip (Ciprofloxacin) resistance also means resistance to nalidixic acid (Nal).

| | Internal ID | Strain | Origin | Sample | Place of origin | Year | Serotype | ST (PFGE) | Phylo group | Resistance phenotype | References |
|---|---|---|---|---|---|---|---|---|---|---|---|
| 1 | SIB1 | 255/16 | broiler | neckskin | Tolna | 2016 | 7:r:1,5 | 32 (B2) | | Amp-Cip-Sul-Tet | this study |
| 2 | SIB2 | SM-1071 | broiler | faeces | Veszprém | 2016 | 7:r:1,5 | 32 (C) | | Amp-Ctx | this study |
| 3 | SIB3 | 263/16 | broiler | neckskin | Tolna | 2016 | 7:r:1,5 | 32 (B2) | | Amp-Cip-Sul-Tet | this study |
| 4 | SIB4 | 343/16 | broiler | neckskin | Tolna | 2016 | 7:r:1,5 | 32 (B2) | | Amp-Cip-Sul-Tet | this study |
| 5 | SIB5 | SM-2416/11 | broiler | faeces | Borsod-Abaúj-Zemplén | 2011 | 7:r:1,5 | 32 (B2) | | Amp-Cip-Sul-Tet | Szmolka et al., 2018 |
| 6 | SIB6 | SM-1174 | broiler | faeces | | 2016 | 7:r:1,5 | 7082 (C) | | Amp-Nal-Sul-Tet | this study |
| 7 | SIB7 | SM-1331 | broiler | faeces | Baranya | 2016 | 7:r:1,5 | 32 (B2) | | Amp-Cip-Sul-Tet | this study |
| 8 | SIB8 | SM-1124/11 | broiler | faeces | Bács-Kiskun | 2011 | 7:r:1,5 | 32 (B1) | | Nal-Sul-Tet | Szmolka et al., 2018 |
| 9 | SIB9 | SM-1160/11 | broiler | faeces | Békés | 2011 | 7:r:1,5 | 32 (P1) | | Nal-Sul-Tet | Szmolka et al., 2018 |
| 10 | SIB10 | SM-1584/11 | broiler | faeces | Somogy | 2011 | 7:r:1,5 | 32 (M1) | | Nal-Sul-Tet | Szmolka et al., 2018 |
| 11 | SIB11 | SM-2041/11 | broiler | faeces | Békés | 2011 | 7:r:1,5 | 32 (B2) | | Nal-Sul-Tet | Szmolka et al., 2018 |
| 12 | SIB12 | SM-630/11 | broiler | faeces | Vas | 2011 | 7:r:1,5 | 32 (O1) | | Nal-Sul-Tet | Szmolka et al., 2018 |
| 13 | SIB13 | SM-635/11 | broiler | faeces | Borsod-Abaúj-Zemplén | 2011 | 7:r:1,5 | 32 (B2) | | Nal-Sul-Tet | Szmolka et al., 2018 |
| 14 | SIB14 | SM-981/11 | broiler | faeces | Győr-Moson-Sopron | 2011 | 7:r:1,5 | 32 (B2) | | Nal-Sul-Tet | Szmolka et al., 2018 |
| 15 | SIB15 | SM-1420/12 | broiler | faeces | Szabolcs-Szatmár-Bereg | 2012 | 7:r:1,5 | 32 (B2) | | Nal-Sul-Tet | Szmolka et al., 2018 |
| 16 | SIB16 | SM-1896/12 | broiler | faeces | Komárom-Esztergom | 2012 | 7:r:1,5 | 32 (B2) | | Nal-Sul-Tet | Szmolka et al., 2018 |
| 17 | SIB17 | SM-2370/12 | broiler | faeces | Győr-Moson-Sopron | 2012 | 7:r:1,5 | 32 (B2) | | Nal-Sul-Tet | Szmolka et al., 2018 |
| 18 | SIB18 | SM-2818/12 | broiler | faeces | Pest | 2012 | 7:r:1,5 | 32 (B2) | | Nal-Sul-Tet | Szmolka et al., 2018 |
| 19 | SIB19 | SM-3364/12 | broiler | faeces | Komárom-Esztergom | 2012 | 7:r:1,5 | 32 (B2) | | Nal-Sul-Tet | Szmolka et al., 2018 |
| 20 | SIB20 | SM-98/12 | broiler | faeces | Vas | 2012 | 7:r:1,5 | 32 (N1) | | Nal-Sul-Tet | Szmolka et al., 2018 |
| 21 | SIB21 | SM-514/13 | broiler | faeces | Veszprém | 2013 | 7:r:1,5 | 32 (B2) | | Nal-Sul-Tet | Szmolka et al., 2018 |
| 22 | SIB22 | SM-521/13 | broiler | faeces | Győr-Moson-Sopron | 2013 | 7:r:1,5 | 32 (B2) | | Nal-Sul-Tet | Szmolka et al., 2018 |
| 23 | SIB23 | SM-750/13 | broiler | faeces | Hajdú-Bihar | 2013 | 7:r:1,5 | 32 (B2) | | Nal-Sul-Tet | Szmolka et al., 2018 |
| 24 | SIB24 | SM-799/13 | broiler | faeces | Somogy | 2013 | 7:r:1,5 | 32 (L1) | | Nal-Sul-Tet | Szmolka et al., 2018 |

| | | | | | | | | | | | |
|---|---|---|---|---|---|---|---|---|---|---|---|
| 25 | | SIB25 | SM-814/13 | broiler | faeces | Baranya | 2013 | 7:r:1,5 | 32 (B2) | Nal-Sul-Tet | Szmolka et al., 2018 |
| 26 | • | SI1/1 | 17991-1S | broiler | caecum | Farm 1 | 2018 | 7:r:1,5 | 32 (B2) | Amp-Cip-Sul-Tet | this study |
| 27 | • | SI4/1 | 18077-1S | broiler | caecum | Farm 2 | 2018 | 7:r:1,5 | 32 (C) | Nal-Sul-Tet | this study |
| 28 | • | SI9/1 | 19740-1S | broiler | caecum | Farm 3 | 2018 | 7:r:1,5 | 32 (B2) | Nal-Sul-Tet | this study |
| 29 | • | SI10/1 | 19711-1S | broiler | caecum | Farm 4 | 2018 | 7:r:1,5 | 32 (B2) | Amp-Nal-Sul-Tet | this study |
| 30 | • | SI12/1 | 21191-1S | broiler | caecum | Farm 5 | 2018 | 7:r:1,5 | 32 (B2) | Nal-Sul-Tet | this study |
| 31 | • | SI15/2 | 23004-2S | broiler | caecum | Farm 6 | 2018 | 7:r:1,5 | 32 (B2) | Nal-Sul-Tet | this study |
| 1 | | SIH1 | 711/12 | human | | Veszprém | 2012 | 7:r:1,5 | 32 (A5) | Amp-Chl-Sul-Tet-Tmp | Szmolka et al., 2018 |
| 2 | | SIH2 | 584/12 | human | | Budapest | 2012 | 7:r:1,5 | 32 (B1) | Amp-Cip-Sul-Tet | Szmolka et al., 2018 |
| 3 | | SIH3 | 8875/13 | human | faeces | Baranya | 2013 | 7:r:1,5 | 32 (B1) | Amp-Nal-Sul-Tet | Szmolka et al., 2018 |
| 4 | | SIH4 | 599/12 | human | | Pest | 2012 | 7:r:1,5 | 32 (A1) | Amp-Sul-Tmp | Szmolka et al., 2018 |
| 5 | | SIH5 | 9314/13 | human | faeces | Baranya | 2013 | 7:r:1,5 | 32 (M1) | Amp-Sul-Tmp | Szmolka et al., 2018 |
| 6 | | SIH6 | 750/11 | human | | Budapest | 2011 | 7:r:1,5 | 32 (B2) | Nal-Sul-Tet | Szmolka et al., 2018 |
| 7 | | SIH7 | 760/11 | human | | Somogy | 2011 | 7:r:1,5 | 32 (B2) | Nal-Sul-Tet | Szmolka et al., 2018 |
| 8 | | SIH8 | 772/11 | human | | Tolna | 2011 | 7:r:1,5 | 32 (B2) | Nal-Sul-Tet | Szmolka et al., 2018 |
| 9 | | SIH9 | 588/12 | human | | Pest | 2012 | 7:r:1,5 | 32 (A1) | Nal-Sul-Tet | Szmolka et al., 2018 |
| 10 | | SIH10 | 552/12 | human | | Pest | 2012 | 7:r:1,5 | 32 (B1) | Nal-Sul-Tet | Szmolka et al., 2018 |
| 11 | | SIH11 | 589/12 | human | | Nógrád | 2012 | 7:r:1,5 | 32 (B1) | Nal-Sul-Tet | Szmolka et al., 2018 |
| 12 | | SIH12 | 611/12 | human | | Somogy | 2012 | 7:r:1,5 | 32 (B1) | Nal-Sul-Tet | Szmolka et al., 2018 |
| 13 | | SIH13 | 560/12 | human | | Somogy | 2012 | 7:r:1,5 | 32 (B2) | Nal-Sul-Tet | Szmolka et al., 2018 |
| 14 | | SIH14 | 538/12 | human | | Baranya | 2012 | 7:r:1,5 | 32 (B2) | Nal-Sul-Tet | Szmolka et al., 2018 |
| 15 | | SIH15 | 511/12 | human | | Budapest | 2012 | 7:r:1,5 | 32 (B2) | Nal-Sul-Tet | Szmolka et al., 2018 |
| 16 | | SIH16 | 690/12 | human | ascites | Pest | 2012 | 7:r:1,5 | 32 (B2) | Nal-Sul-Tet | Szmolka et al., 2018 |
| 17 | | SIH17 | 718/12 | human | | Somogy | 2012 | 7:r:1,5 | 32 (B2) | Nal-Sul-Tet | Szmolka et al., 2018 |
| 18 | | SIH18 | 613/12 | human | | Győr | 2012 | 7:r:1,5 | 32 (B2) | Nal-Sul-Tet | Szmolka et al., 2018 |
| 19 | | SIH19 | 606/12 | human | | Baranya | 2012 | 7:r:1,5 | 32 (B2) | Nal-Sul-Tet | Szmolka et al., 2018 |
| 20 | | SIH20 | 1818/13 | human | faeces | Veszprém | 2013 | 7:r:1,5 | 32 (B2) | Nal-Sul-Tet | Szmolka et al., 2018 |
| 21 | | SIH21 | 2102/13 | human | hemoculture | Pest | 2013 | 7:r:1,5 | 7081 (B2) | Nal-Sul-Tet | Szmolka et al., 2018 |
| 22 | | SIH22 | 4281/13 | human | faeces | Pest | 2013 | 7:r:1,5 | 32 (B2) | Nal-Sul-Tet | Szmolka et al., 2018 |

| | | | | | | | | | | | |
|---|---|---|---|---|---|---|---|---|---|---|---|
| 23 | SIH23 | 4285/13 | human | hemoculture | Pest | 2013 | 7:r:1,5 | 32 (B2) | | Nal-Sul-Tet | Szmolka et al., 2018 |
| 24 | SIH24 | 5227/13 | human | faeces | Hajdú-Bihar | 2013 | 7:r:1,5 | 32 (B2) | | Nal-Sul-Tet | Szmolka et al., 2018 |
| 25 | SIH25 | 6389/13 | human | faeces | Pest | 2013 | 7:r:1,5 | 32 (B2) | | Nal-Sul-Tet | Szmolka et al., 2018 |
| 1 | Ec1 | PA1-1Ec Ctx | broiler | faeces | Fót I (chicken 1) | 2013 | O159:H20 | 2040 | A | Amp-Chl-Ctx-Gen-Sul-Tet | this study |
| 2 | Ec2 | PA11-2Ec Nal | broiler | faeces | Fót I (chicken 11) | 2013 | ONt:H34 | 354 | F | Amp-Gen-Cip-Sul-Tet-Tmp | this study |
| 3 | Ec3 | PC3-1Ec Ctx | broiler | faeces | Galgamácsa (chicken 3) | 2013 | O8:H9 | 23 | C | Amp-Chl-Ctx-Gen-Sul-Tet | this study |
| 4 | Ec4 | PB5-1Ec Ctx | broiler | faeces | Fót II (chicken 5) | 2013 | O83:H42 | 1485 | F | Amp-Ctx-Cip-Sul-Tet-Tmp | this study |
| 5 | Ec5 | PB15-2Ec Nal | broiler | faeces | Fót II (chicken 15) | 2013 | O166:H25 | 10 | A | Amp-Ctx-Gen-Nal-Sul-Tet | this study |
| 6 | Ec6 | PC10-2Ec Ctx | broiler | faeces | Galgamácsa (chicken 10) | 2013 | O10:H9 | 115 | D | Amp-Cip-Ctx-Sul-Tet-Tmp | this study |
| 7 | Ec7 | PB14-1Ec Ctx | broiler | faeces | Fót II (chicken 14) | 2013 | O17/O44/O77:HNt | 1158 | F | Amp-Ctx-Nal | this study |
| 8 | Ec8 | PC15-1Ec Ctx | broiler | faeces | Galgamácsa (chicken 15) | 2013 | O50/O2:H1 | 429 | B2 | Amp-Ctx-Sul-Tet | this study |
| 9 | Ec9 | PC5-3Ec Nal | broiler | faeces | Galgamácsa (chicken 5) | 2013 | O8:H7 | 1642 | B1 | Amp-Cip-Gen-Sul-Tmp | this study |
| 10 | Ec10 | PA5-1Ec Nal | broiler | faeces | Fót I (chicken 5) | 2013 | O102:H23 | 224 | B1 | Amp-Chl-Cip-Sul-Tet-Tmp | this study |
| 11 | Ec11 | PB6-1Ec Nal | broiler | faeces | Fót II (chicken 6) | 2013 | ONt:H40 | 398 | A | Amp-Cip-Tet | this study |
| 12 | Ec12 | PC4-1Ec Nal | broiler | faeces | Galgamácsa (chicken 4) | 2013 | ONt:H40 | 398 | A | Amp-Cip-Sul-Tet-Tmp | this study |
| 13 | Ec13 | PA1-3Ec Nal | broiler | faeces | Fót I (chicken 1) | 2013 | O11:H4 | 117 | F | Chl-Cip-Gen-Sul-Tet-Tmp | this study |
| 14 | Ec14 | PA10-1Ec Ctx | broiler | faeces | Fót I (chicken 10) | 2013 | O8:H9 | 23 | C | Amp-Ctx | this study |
| 15 | Ec15 | PB8-2Ec Ctx | broiler | faeces | Fót II (chicken 8) | 2013 | O15:H6 | 2309 | D | Amp-Ctx-Tet | this study |
| 16 | Ec16 | PC1-1Ec Ctx | broiler | faeces | Galgamácsa (chicken 1) | 2013 | O11:H43 | 7705 | B1 | Amp-Ctx | this study |
| 17 | Ec17 | PC8-3Ec Nal | broiler | faeces | Galgamácsa (chicken 8) | 2013 | O9:H21 | 155 | B1 | Amp-Nal | this study |
| 18 | Ec18 | PB2-1Ec Nal | broiler | faeces | Fót II (chicken 2) | 2013 | O159:H21 | 641 | B1 | Amp-Cip | this study |
| 19 | Ec19 | PC6-1Ec Nal | broiler | faeces | Galgamácsa (chicken 6) | 2013 | O18:H7 | 351 | B1 | Amp-Cip-Sul-Tmp | this study |
| 20 | Ec20 | PA2-2Ec Nal | broiler | faeces | Fót I (chicken 2) | 2013 | O9:H21 | 155 | B1 | Amp-Cip-Tet | this study |
| 21 | Ec21 | PA3-2Ec Nal | broiler | faeces | Fót I (chicken 3) | 2013 | O154:H10 | 533 | B1 | Cip-Tet | this study |
| 22 | Ec22 | PB7-1Ec Nal | broiler | faeces | Fót II (chicken 7) | 2013 | O82:H21 | 1800 | B1 | Cip-Tet | this study |

| | | | | | | | | | | | | |
|---|---|---|---|---|---|---|---|---|---|---|---|---|
| 1 | • | Ec1/1 | 17991-1Ec | broiler | caecum | Farm 1 | 2018 | O9:H9 | 2179 | B1 | Amp-Nal-Tet | this study |
| 2 | • | Ec1/2 | 17991-2Ec | broiler | caecum | Farm 1 | 2018 | O9:H19 | 155 | B1 | Amp-Cip-Tet | this study |
| 3 | • | Ec1/5 | 17991-5Ec | broiler | caecum | Farm 1 | 2018 | O61:H51 | 155 | B1 | Amp-Chl-Cip-Sul-Tet-Tmp | this study |
| 4 | • | Ec1/7 | 17991-7Ec | broiler | caecum | Farm 1 | 2018 | ONt:H14 | 1249 | A | Cip-Sul-Tet-Tmp | this study |
| 5 | • | Ec4/1 | 18077-1Ec | broiler | caecum | Farm 2 | 2018 | O29:H10 | 1720 | B1 | Amp | this study |
| 6 | • | Ec4/2 | 18077-2Ec | broiler | caecum | Farm 2 | 2018 | O171:H4 | 117 | F | Sul | this study |
| 7 | • | Ec4/3 | 18077-3Ec | broiler | caecum | Farm 2 | 2018 | O100:H25 | 683 | B1 | Amp-Chl-Tet | this study |
| 8 | • | Ec4/5 | 18077-5Ec | broiler | caecum | Farm 2 | 2018 | O9:H9 | 162 | B1 | Cip-Sul | this study |
| 9 | • | Ec4/9 | 18077-9Ec | broiler | caecum | Farm 2 | 2018 | O9:H4 | 46 | A | Nal | this study |
| 10 | • | Ec9/2 | 19740-2Ec | broiler | caecum | Farm 3 | 2018 | O26:H34 | 752 | A | Amp-Chl-Nal-Tet | this study |
| 11 | • | Ec9/3 | 19740-3Ec | broiler | caecum | Farm 3 | 2018 | O75:H42 | 8701 | A | Amp-Cip-Sul-Tet-Tmp | this study |
| 12 | • | Ec9/4 | 19740-4Ec | broiler | caecum | Farm 3 | 2018 | O88:H12 | 10 | A | Amp-Ctx | this study |
| 13 | • | Ec9/6 | 19740-6Ec | broiler | caecum | Farm 3 | 2018 | O160:H4 | 117 | F | Amp-Cip-Ctx | this study |
| 14 | • | Ec9/7 | 19740-7Ec | broiler | caecum | Farm 3 | 2018 | O18:H49 | 212 | B1 | Amp-Chl-Cip-Sul-Tet | this study |
| 15 | • | Ec10/2 | 19711-2Ec | broiler | caecum | Farm 4 | 2018 | ONt:H4 | 665 | A | Cip | this study |
| 16 | • | Ec10/6 | 19711-6Ec | broiler | caecum | Farm 4 | 2018 | ONt:H48 | 10 | A | Amp-Nal | this study |
| 17 | • | Ec10/7 | 19711-7Ec | broiler | caecum | Farm 4 | 2018 | O69:H38 | 3107 | A | Amp-Sul-Tet | this study |
| 18 | • | Ec10/9 | 19711-9Ec | broiler | caecum | Farm 4 | 2018 | O21:H45 | 616 | B1 | Amp-Sul-Tet-Tmp | this study |
| 19 | • | Ec12/1 | 21191-1Ec | broiler | caecum | Farm 5 | 2018 | O8:H9 | 8702 | A | Amp-Cip-Tet | this study |
| 20 | • | Ec12/3 | 21191-3Ec | broiler | caecum | Farm 5 | 2018 | O7:H31 | 58 | B1 | Cip | this study |
| 21 | • | Ec12/4 | 21191-4Ec | broiler | caecum | Farm 5 | 2018 | O4:H4 | 10 | A | - | this study |
| 22 | • | Ec12/5 | 21191-5Ec | broiler | caecum | Farm 5 | 2018 | O4:H4 | 10 | A | Amp | this study |
| 23 | • | Ec12/6 | 21191-6Ec | broiler | caecum | Farm 5 | 2018 | O117:H45 | 616 | B1 | Nal | this study |
| 24 | • | Ec15/1 | 23004-1Ec | broiler | caecum | Farm 6 | 2018 | O3:H10 | 118 | E | Nal | this study |
| 25 | • | Ec15/2 | 23004-2Ec | broiler | caecum | Farm 6 | 2018 | O3:H10 | 118 | E | Cip | this study |
| 26 | • | Ec15/3 | 23004-3Ec | broiler | caecum | Farm 6 | 2018 | O16:H48 | 10 | A | Cip | this study |
| 27 | • | Ec15/10 | 23004-10Ec | broiler | caecum | Farm 6 | 2018 | ONt:H52 | 3941 | A | Amp-Chl-Cip-Sul-Tet | this study |
| 28 | | IntEC1/1 | 1208/vb-1Ec | day old chick | caecum | Hegyháthodász | 2018 | O78:H4 | 117 | F | Cip | this study |

| | | | | | | | | | | | |
|---|---|---|---|---|---|---|---|---|---|---|---|
| 29 | IntEC2/1 | 1211/vb-1Ec | day old chick | caecum | Adásztevel | 2018 | O23:H16 | 453 | B1 | Amp-Cip-Chl-Sul-Tmp | this study |
| 30 | IntEC3/1 | 1232/vb-1Ec | day old chick | caecum | Hegyháthodász | 2018 | O9:H19 | 162 | B1 | Nal | this study |
| 31 | IntEC3/8 | 1232/vb-8Ec | day old chick | caecum | Hegyháthodász | 2018 | O50/O2:H5 | 355 | B2 | Nal | this study |
| 32 | IntEC3/9 | 1232/vb-9Ec | day old chick | caecum | Hegyháthodász | 2018 | O51:H37 | 297 | B1 | Amp-Cip-Tet-Tmp | this study |
| 33 | IntEC4/1 | 1437/vb-1Ec | day old chick | caecum | Völcsej | 2018 | O51:H10 | 10088 | A | - | this study |
| 34 | IntEC4/5 | 1437/vb-5Ec | day old chick | caecum | Völcsej | 2018 | O50/O2:H5 | 95 | B2 | Nal-Sul | this study |
| 35 | IntEC5/1 | 3921F/vb-1Ec | day old chick | caecum | Földeák | 2018 | O8:H25 | 58 | B1 | Amp-Cip-Tet | this study |
| 36 | IntEC6/1 | 3922F/vb-1Ec | day old chick | caecum | Földeák | 2018 | O166:H15 | 349 | D | - | this study |
| 37 | IntEC6/5 | 3922F/vb-5Ec | day old chick | caecum | Földeák | 2018 | O8:H25 | 58 | B1 | Amp-Nal-Tet | this study |
| 38 | IntEC9/1 | 403/vb-1Ec | day old chick | caecum | Békés | 2018 | O5:H21 | 162 | B1 | Amp-Cip-Sul-Tet-Tmp | this study |
| 39 | IntEC9/3 | 403/vb-3Ec | day old chick | caecum | Békés | 2018 | O5:H40 | 93 | A | Nal | this study |
| 40 | IntEC11/1 | 1588/vb-1Ec | day old chick | caecum | Pakod | 2018 | O114:H4 | 117 | F | - | this study |
| 41 | IntEC11/6 | 1588/vb-6Ec | day old chick | caecum | Pakod | 2018 | O78:H4 | 117 | F | Amp-Sul-Tmp | this study |
| 42 | IntEC12/1 | 1589/vb-1Ec | day old chick | caecum | Kisvásárhely | 2018 | O88:H10 | 162 | B1 | Amp-Cip-Sul-Tmp | this study |
| 43 | IntEC12/3 | 1589/vb-3Ec | day old chick | caecum | Kisvásárhely | 2018 | O8:H9 | 90 | C | Amp-Cip-Sul | this study |
| 1 | ExPEC1/1 | 1208/csv-1Ec | day old chick | bone marrow | Hegyháthodász | 2018 | O78:H4 | 117 | F | Cip | this study |
| 2 | ExPEC1/4 | 1208/csv-4Ec | day old chick | bone marrow | Hegyháthodász | 2018 | O23:H16 | 453 | B1 | Amp-Cip-Chl-Sul-Tmp | this study |
| 3 | ExPEC2/1 | 1211/csv-1Ec | day old chick | bone marrow | Adásztevel | 2018 | O23:H16 | 453 | B1 | Amp-Cip-Chl-Sul-Tmp | this study |
| 4 | ExPEC3/1 | 1232/csv-1Ec | day old chick | bone marrow | Hegyháthodász | 2018 | O50/O2:H5 | 355 | B2 | Nal | this study |
| 5 | ExPEC4/1 | 1437/csv-1Ec | day old chick | bone marrow | Völcsej | 2018 | O78:H4 | 117 | F | - | this study |
| 6 | ExPEC5/1 | 3921F/csv-1Ec | day old chick | bone marrow | Földeák | 2018 | O5:H10 | 93 | A | - | this study |
| 7 | ExPEC6/1 | 3922F/csv-1Ec | day old chick | bone marrow | Földeák | 2018 | O88:H8 | 101 | B1 | Nal | this study |

| | | | | | | | | | | | |
|---|---|---|---|---|---|---|---|---|---|---|---|
| 8 | ExPEC9/1 | 403/csv-1Ec | day old chick | bone marrow | Békés | 2018 | O5:H40 | 93 | A | Nal | this study |
| 9 | ExPEC11/2 | 1588/csv-2Ec | day old chick | bone marrow | Pakod | 2018 | O8:H19 | 88 | C | Amp-Chl-Sul-Tet-Tmp | this study |
| 10 | ExPEC12/1 | 1589/csv-1Ec | day old chick | bone marrow | Kisvásárhely | 2018 | O88:H10 | 162 | B1 | Amp-Cip-Sul-Tmp | this study |
| 11 | ExPEC227 | 411/2b | day old chick | bone marrow | Pölöske | 1998-1999 | O15:H10 | 93 | A | Nal-Tet | Tóth et al., 2012 |
| 12 | ExPEC229 | K-II-8 114 | day old chick | bone marrow | Devecser | 1998-1999 | O100:H32 | 10 | A | Amp-Sul | Tóth et al., 2012 |
| 13 | ExPEC235 | K-II-35 163 | day old chick | bone marrow | Szombathely | 1998-1999 | ONt:H28 | 348 | B1 | Amp-Nal-Tet | Tóth et al., 2012 |
| 14 | ExPEC241 | K-II-49 177 | day old chick | bone marrow | Vasvár | 1998-1999 | O78:H12 | 88 | C | Amp-Nal-Sul-Tet-Tmp | Tóth et al., 2012 |
| 15 | ExPEC243 | 378/4a | day old chick | bone marrow | Szombathely | 1998-1999 | O78:H12 | 88 | C | Amp-Nal-Sul-Tet-Tmp | Tóth et al., 2012 |
| 16 | ExPEC249 | 378/5a | day old chick | bone marrow | Szombathely | 1998-1999 | O115:H4 | 117 | F | Amp-Sul-Tet-Tmp | Tóth et al., 2012 |
| 17 | ExPEC256 | K-II-46 174 | day old chick | bone marrow | Vasvár | 1998-1999 | O115:H4 | 117 | F | Amp-Sul-Tet-Tmp | Tóth et al., 2012 |
| 18 | ExPEC292 | K-II-45 173 | day old chick | bone marrow | Vasvár | 1998-1999 | O6:H16 | 373 | A | Nal-Tet | Tóth et al., 2012 |
| 19 | ExPEC699 | 390/3b | day old chick | bone marrow | Simaság | 1999-2000 | O120:H4 | 428 | B2 | Nal-Tet | this study |
| 20 | ExPEC712 | 389/3a | day old chick | bone marrow | Simaság | 1999-2000 | O53:H4 | 117 | F | Amp-Nal-Sul-Tet-Tmp | this study |
| 21 | ExPEC718 | 381/2a | day old chick | bone marrow | Rábaipoly | 1999-2000 | O53:H4 | 117 | F | Amp-Nal-Sul-Tet-Tmp | this study |
| 22 | ExPEC726 | 379/3a | day old chick | bone marrow | Szombathely | 1999-2000 | O182:H7 | 1148 | B1 | Tet | this study |
| 23 | ExPEC735 | 381/1a | day old chick | bone marrow | Rábaipoly | 1999-2000 | O88:H8 | 101 | B1 | Gen-Nal-Sul-Tet | this study |
| 24 | ExPEC736 | K-II-21 | day old chick | bone marrow | Devecser | 1999-2000 | O88:H8 | 101 | B1 | Gen-Nal-Sul-Tet | this study |
| 25 | ExPEC739 | K-II-27 | day old chick | bone marrow | Devecser | 1999-2000 | O89:H10 | 10 | A | Nal | this study |
