## Supplemental Table 2 for "Comparative genomics of emerging lineages and mobile resistomes of contemporary broiler strains of *Salmonella* Infantis and *E. coli*"

**Table S2. Sero- and phylogroups of *E. coli* strains isolated from different broiler sources.**

| | O-type | H-type | Phylogroup | ST | Strain no. | |
|---|---|---|---|---|---|---|
| 1 | O3 | H10 | E | 118 | 2 | |
| 2 | O4 | H4 | A | 10 | 2 | |
| 3 | O5 | H10 | A | 93 | 1 | |
| | | H21 | B1 | 162 | 1 | |
| | | H40 | A | 93 | 1 | |
| | | | A | 93 | 1 | **4** |
| 4 | O6 | H16 | A | 373 | 1 | |
| 5 | O7 | H31 | B1 | 58 | 1 | |
| 6 | O8 | H7 | B1 | 1642 | 1 | |
| | | H9 | A | nt | 1 | |
| | | | C | 90 | 1 | |
| | | | C | 23 | 2 | |
| | | H19 | C | 88 | 1 | |
| | | H25 | B1 | 58 | 2 | **8** |
| 7 | O9 | H4 | A | 46 | 1 | |
| | | H9 | B1 | 162 | 1 | |
| | | | B1 | 2179 | 1 | |
| | | H19 | B1 | 155 | 1 | |
| | | | B1 | 162 | 1 | |
| | | H21 | B1 | 155 | 2 | **7** |
| 8 | O10 | H9 | D | 115 | 1 | |
| 9 | O11 | H4 | F | 117 | 1 | |
| | | H43 | B1 | 7705 | 1 | |
| 10 | O15 | H6 | D | 2309 | 1 | |
| | | H10 | A | 93 | 1 | |
| 11 | O16 | H48 | A | 10 | 1 | |
| 12 | O17/O44/O77 | nt | F | 1158 | 1 | |
| 13 | O18 | H7 | B1 | 351 | 1 | |
| | | H49 | B1 | 212 | 1 | |
| 14 | O21 | H45 | B1 | 616 | 1 | |
| 15 | O23 | H16 | B1 | 453 | 2 | |
| | | | B1 | 453 | 1 | |
| 16 | O26 | H34 | A | 752 | 1 | |
| 17 | O29 | H10 | B1 | 1720 | 1 | |
| 18 | O50/O2 | H1 | B2 | 429 | 1 | |
| | | H5 | B2 | 95 | 1 | |
| | | | B2 | 355 | 1 | |
| | | | B2 | 355 | 1 | **4** |

**SAMPLE SOURCE**

- CAECUM
- BONE MARROW
- FAECES

| | | | | | | |
|---|---|---|---|---|---|---|
| 19 | O51 | H10 | A | nt | 1 | |
| | | H37 | B1 | 297 | 1 | |
| 20 | O53 | H4 | F | 117 | 2 | |
| 21 | O61 | H51 | B1 | 155 | 1 | |
| 22 | O69 | H38 | A | 3107 | 1 | |
| 23 | O75 | H42 | A | 8701 | 1 | |
| 24 | O78 | H4 | F | 117 | 2 | |
| | | | F | 117 | 2 | |
| | | H12 | C | 88 | 2 | **6** |
| 25 | O82 | H21 | B1 | 1800 | 1 | |
| 26 | O83 | H42 | F | 1485 | 1 | |
| 27 | O88 | H8 | B1 | 101 | 3 | |
| | | H10 | B1 | 162 | 1 | |
| | | | B1 | 162 | 1 | |
| | | H12 | A | 10 | 1 | **6** |
| 28 | O89 | H10 | A | 10 | 1 | |
| 29 | O100 | H25 | B1 | 683 | 1 | |
| | | H32 | A | 10 | 1 | |
| 30 | O102 | H23 | B1 | 224 | 1 | |
| 31 | O114 | H4 | F | 117 | 1 | |
| 32 | O115 | H4 | F | 117 | 2 | |
| 33 | O117 | H45 | B1 | 616 | 1 | |
| 34 | O120 | H4 | B2 | 428 | 1 | |
| 35 | O154 | H10 | B1 | 533 | 1 | |
| 36 | O159 | H20 | A | 2040 | 1 | |
| | | H21 | B1 | 641 | 1 | |
| 37 | O160 | H4 | F | 117 | 1 | |
| 38 | O166 | H15 | D | 349 | 1 | |
| | | H25 | A | 10 | 1 | |
| 39 | O171 | H4 | F | 117 | 1 | |
| 40 | O182 | H7 | B1 | 1148 | 1 | |
| | nt | H4 | A | 665 | 1 | |
| | nt | H14 | A | 1249 | 1 | |
| | nt | H28 | B1 | 348 | 1 | |
| | nt | H34 | F | 354 | 1 | |
| | nt | H40 | A | 398 | 2 | |
| | nt | H48 | A | 10 | 1 | |
| | nt | H52 | A | 3941 | 1 | **8** |
| | | | | **90** | | |
